# Structural analysis of the lncRNA SChLAP1 reveals protein binding interfaces and a conformationally heterogenous retroviral insertion

**DOI:** 10.1101/2021.10.21.465303

**Authors:** James P. Falese, Emily J. McFadden, Christopher A. D’Inzeo, Amanda E. Hargrove

## Abstract

The lncRNA Second Chromosome Locus Associated with Prostate 1 (SChLAP1) was previously identified as a predictive biomarker and potential driver of aggressive prostate cancer. Recent work suggests that SChLAP1 may bind the SWI/SNF chromatin remodeling complex to promote prostate cancer metastasis, though the exact role of SWI/SNF recognition is debated. To date, there are no detailed biochemical studies of *apo* SChLAP1 or SChLAP1:protein complexes. Herein, we report the first secondary structure model of SChLAP1 using SHAPE-MaP *in* vitro, *in cellulo*, and *ex cellulo* (protein-free). Comparison of the *ex cellulo* and *in cellulo* data via ΔSHAPE identified putative protein binding sites within SChLAP1. In addition, we identified primate conserved exons of SChLAP1 as well as regions that appear to have been incorporated through retroviral insertion. In particular, we characterized a complex structural landscape in a protein binding region at the 3′—end of SChLAP1 derived from a THE1B-type retroviral insertion, suggesting a role for an exapted RNA structure in SChLAP1:protein recognition and prostate cancer progression. This work lays the foundation for future efforts to selectively target and disrupt the SChLAP1:protein interface and to develop new therapeutic avenues in prostate cancer treatment.

## INTRODUCTION

Prostate cancer is one of the most commonly diagnosed cancers among American men, alone accounting for 27% of all cancer diagnoses in men, and it is the second leading cause of cancer death in American men (Siegel et al. 2022). While several treatment options are available for prostate cancer, including prostatectomy, radiation-based therapies, chemotherapy, immunotherapy, and hormone deprivation therapies (PDQ® Adult Treatment Editorial Board, https://www.cancer.gov/types/prostate/patient/prostate-treatment-pdq), none of these treatments provide a cure for prostate cancer. Additionally, the subsequent emergence of metastatic prostate cancer, also known as aggressive prostate cancer, in a subset of patients rapidly results in treatment resistance and a significantly bleaker prognosis (Siegel et al. 2022). Given the high incidence of prostate cancer and the correspondingly large clinical burden of metastatic prostate cancer, there is an urgent and unmet need for specific therapeutic strategies that target molecular drivers of aggressive prostate cancer.

Noncoding RNA remains an underexplored area for therapeutic development, particularly via small molecule-based therapies (Warner et al. 2018; Costales et al. 2020; Falese et al. 2021). Long non-coding RNAs (lncRNAs), generally defined as non-translated transcripts ≥200 nucleotides in length, are often differentially expressed throughout developmental stages, tissue types, and disease states (Mattick et al. 2023). Following the ENCODE project and identification of thousands of new lncRNAs (Consortium 2012; Djebali et al. 2012), additional work, particularly from the FANTOM consortium, began annotating these transcripts and found biochemical indices of function historically ascribed solely to proteins (Carninci et al. 2005; Hon et al. 2017). Although there are over 100,000 identified lncRNA (Mattick et al. 2023), only ∼20,000 of these have been implicated in biological function (Hon et al. 2017), and an even smaller portion of these have been biochemically characterized for either housekeeping or disease/cancer-associated functions.

The lncRNA Second Chromosome Locus Associated with Prostate-1 (SChLAP1) is a prime example of the incongruity between the identification of cancer-associated lncRNAs and characterization of their biochemical function. In 2011, Prensner et al. identified the transcript Prostate Cancer Associated Transcript 114 (PCAT-114; later renamed SChLAP1) as overexpressed in tumor prostate tissue compared to benign prostate tissue (Prensner et al. 2011). In other work, Gerashchenko et al. determined that patient-derived tumors with high SChLAP1 transcript levels correlated with a Gleason score of 9, where 10 is the highest possible score and most at-risk group (Gerashchenko et al. 2018). Additionally, SChLAP1 levels correlated with high levels of epithelial-to-mesenchymal transition (EMT) markers such as vimentin (VIM), fibronectin (FN1), and matrix metalloproteinase 2 (MMP2), further supporting the correlation between SChLAP1 overexpression and prostate cancer metastasis found in multiple clinical studies (Prensner et al. 2013; Prensner et al. 2014; Mehra et al. 2014; Mehra et al. 2016; Chua et al. 2017; Gerashchenko et al. 2018; Kidd et al. 2021). Furthermore, Mehra et al. determined that approximately 16% of clinically localized prostate cancers in American men exhibit high SChLAP1 transcript levels, suggesting use of SChLAP1 as an early detection biomarker (Mehra et al. 2014). A mechanistic role for SChLAP1 in promoting prostate cancer progression was also supported in other cell-based *in vitro* and *in vivo* experiments. Intracardiac injection of 22Rv1 (human-derived prostate cancer) cells with stable SChLAP1 knockdown into severe combined immunodeficiency mice showed a significant reduction in the number and size of metastases as compared to a non-targeting shRNA negative control. In addition, siRNA-mediated knockdown of SChLAP1 in human prostate cancer cell lines significantly reduced *in vitro* invasion in Boyden Chamber Assays as compared to non-targeting control siRNA and displayed equivalent results as compared to siRNA-mediated knockdown of *EZH2* mRNA (siEZH2), a known promoter of cell proliferation (Prensner et al. 2013).

Furthermore, Prensner et al. showed that overexpression of SChLAP1 in the non-cancerous RWPE-1 (human healthy prostate) cell line via lentiviral transduction transformed the usually non-invasive prostate cells into invasive prostate cells (as assessed by a Boyden chamber assay), supporting SChLAP1 as sufficient for promoting invasion, which was also confirmed in an independent study (Prensner et al. 2013; Raab et al. 2019). From these works, SChLAP1 overexpression was identified as a factor in three hallmarks of carcinogenesis: metastasis, proliferation, and invasion. However, despite the demonstrated biological importance of SChLAP1 in prostate cancer progression, identification of a molecular mechanism has remained elusive. Initial work from Prensner et al. identified a specific interaction between SChLAP1 and the SWItch/Sucrose Non-Fermentable (SWI/SNF) chromatin remodeling complex, which catalyzes ATP-dependent chromatin remodeling through sliding or ejection of nucleosomes from DNA (Prensner et al. 2013). The SWI/SNF complex exhibits mutations in at least one of its component subunits in nearly 25% of all cancers (Roberts and Orkin 2004; Shain and Pollack 2013; Kadoch et al. 2016; Mittal and Roberts 2020). Specifically, Prensner et al. observed SChLAP1-dependent eviction of the SWI/SNF complex from chromatin, resulting in gene expression changes and facilitating an aggressive phenotype (Prensner et al. 2013). However, more recent work has identified broad, potentially non-specific interactions between the SWI/SNF complex and RNA (Cajigas et al. 2015; Raab et al. 2019; Grossi et al. 2020; Skalska et al. 2021). In addition, Raab et al. observed that SWI/SNF remained chromatin-associated regardless of SChLAP1 expression, suggesting that genome-wide eviction of SWI/SNF was not the source of the SChLAP1-induced phenotype and that other protein interactions may be required for SChLAP1-promoted cancer aggression (Raab et al. 2019). Since these studies, several other proteins and/or protein complexes have been proposed to interact with SChLAP1, including 1) Polycomb repressive complex 2 (PRC2), a histone methyltransferase (Huang and Tang 2021), 2) DNA (cytosine-5)-methyltransferase 3A (DNMT3A), a DNA methyltransferase (Huang and Tang 2021), 3) Heterogeneous nuclear ribonucleoprotein L (HNRNPL) (Ji et al. 2019) and 4) Heterogeneous nuclear ribonucleoprotein D0 (HNRNPD) (Du et al. 2021), which are involved in RNA processing. While the interaction between SChLAP1 and HNRNPL was localized to a specific exon within SChLAP1 (Ji et al. 2019), the specific regions of SChLAP1 involved in intermolecular recognition remain undefined.

In summary, despite these identified protein interactions and the established role for SChLAP1 in supporting aggressive prostate cancer, there is little knowledge regarding the RNA features within SChLAP1 that mediate these interactions. Elucidating the sequence- and/or structure-function relationships within SChLAP1 is critical to our understanding of both prostate cancer metastasis and the potential for SChLAP1 as a therapeutic target. Herein, we report first insights into the relationships among SChLAP1 sequence, structure, and function. First, we identified several exons that are highly conserved across all human SChLAP1 isoforms and several non-human primates. Additionally, we identified regions of SChLAP1 that may have been inserted via retroviral insertion during primate evolution. We also generated the first *in vitro* secondary structural model of SChLAP1 isoform 1 *in vitro* via selective 2′-Hydroxyl Acylation Analyzed by Primer Extension and Mutational Profiling (SHAPE-MaP) (Siegfried et al. 2014) and Dimethyl Sulfate Mutational Profiling and sequencing (DMS-MaPseq) (Zubradt et al. 2017). Through our subsequent *in cellulo* and *ex cellulo* SHAPE probing and analyses via ΔSHAPE, we identified putative protein binding regions and significantly unfolded regions across SChLAP1 in cells. Lastly, we identified a complex structural landscape at the 3′—end of SChLAP1 in a region derived from a potential retroviral insertion that additionally corresponds to a putative protein binding region. As the first detailed biochemical and structural analysis of SChLAP1, this work proposes an important functional role of RNA structures within SChLAP1 for protein recognition and ultimately the metastatic phenotype associated with SChLAP1 overexpression in prostate cancer.

## RESULTS

### Evolutionary analysis reveals conserved and potentially functional exons of SChLAP1

Conserved RNA motifs within prokaryotic and eukaryotic transcripts are often indicative of critical function. Thus, we performed a BLAST search of the NCBI RefSeq database to identify potential homologous SChLAP1 sequences in non-human genomes (Zhang et al. 2000; O’Leary et al. 2016). This search revealed several non-human primate (NHP) sequences of SChLAP1, all of which were predicted to be non-coding but have no functional annotation to date (Fig. 1A, Table S1). SChLAP1 sequences were not called in any non-primate species, suggesting that SChLAP1 emerged during primate evolution, consistent with general evidence of greater lncRNA content with increasing organismal complexity and evidence that many lncRNAs are primate specific (Derrien et al 2012; Liu et al. 2013; Mattick et al. 2023). This search also showed evidence of greater conservation of some exons over others; in particular, exons 1, 2, 5, and 7 appeared to be the most conserved across SChLAP1 homologs (Fig. 1A), appearing in 6/9, 9/9, 6/9, and 7/9 sequences, respectively, indicating these exons for a greater likelihood of functional roles. In contrast, exons 3, 4, and 6 were not as commonly observed (Fig. 1A), appearing in 4/9, 1/9, and 4/9 sequences, respectively. We note that no splice isoforms of NHP putative SChLAP1 homologs were identified. The importance of exons 1, 2, 5, and 7 is also indicated by known splice isoforms of SChLAP1 expressed in human tissues. Of the 7 known isoforms in human SChLAP1, all isoforms contain exons 1, 2, and 7, and all but one contain exon 5. In contrast, exons 3, 4, and 6 are observed in 3/7, 2/7, and 4/7 isoforms respectively (not shown). In addition, exons 3 and 4 are not present in the most highly expressed isoforms of SChLAP1 (isoforms 1-3, accounting for 90% of SChLAP1 transcripts in human cells (Prensner et al. 2013)), while exon 6 is only present in one of the top 3 isoforms, suggesting that exons 3, 4, and 6 are dispensable for SChLAP1 activity.

**Figure 1.**
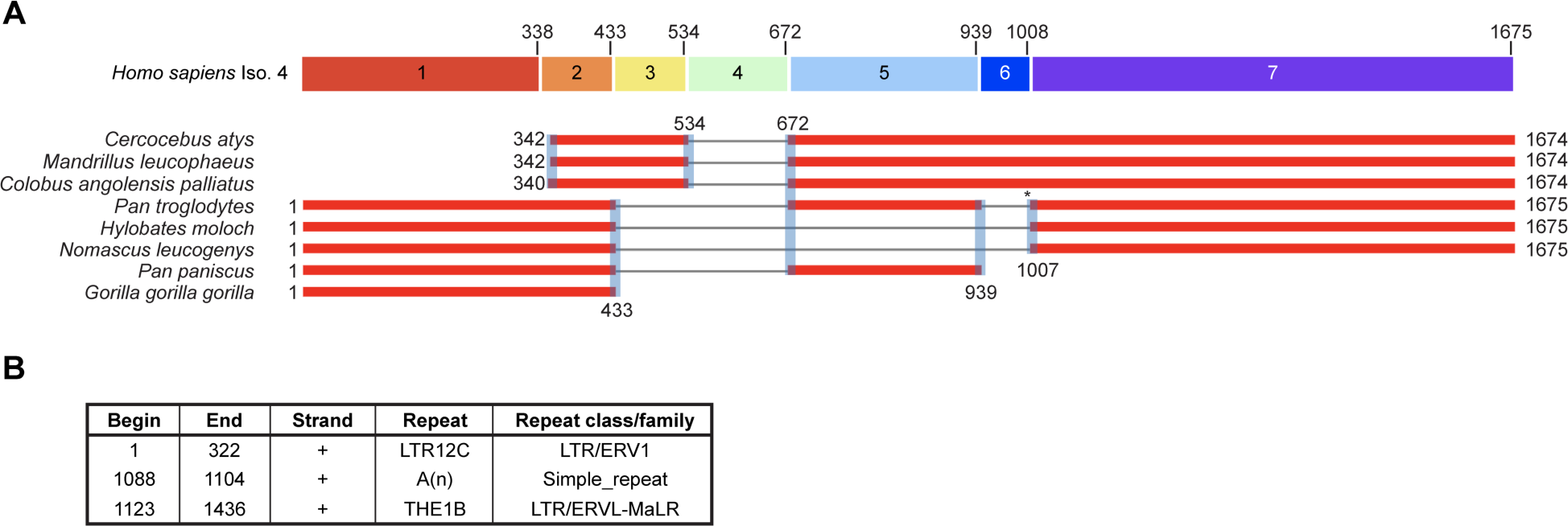
Phylogenetic analysis of SChLAP1. A) BLAST search of isoform 4 of SChLAP1 (containing all possible human exons) against the RefSeq database. Top: Isoform 4 of SChLAP1 with each exon shown for reference. The 3′—positions of each exon in the human SChLAP1 are labeled. Bottom: sequence alignments of all identified SChLAP1 homologs against human SChLAP1 isoform 4. The 5′— and 3′—positions of alignment against human SChLAP1 are noted. All BLAST alignment scores were greater than 200. Asterisk denotes *Pan troglodytes* aligning at nucleotides 1006 rather than 1007. B) Repeatmasker analysis of SChLAP1 Iso 1 (see also Fig. S1).

While our BLAST search uncovered several complete putative SChLAP1 homologs in NHPs, we noted that multiple other genes (both human and NHP, including protein-coding and non-coding genes) aligned with exon 1 of SChLAP1 (not shown); this result suggested that repeat insertions, occurring not only within SChLAP1 but also in other genomic regions, may have resulted in spurious alignment in our BLAST search. These regions (i.e., transposable elements; Tes) are highly promiscuous DNA elements that can replicate independently from the host cell. Tes can become incorporated into coding and non-coding genome sequences and result in the formation of novel functions in lncRNAs, such as protein recognition and/or RNA structures (Johnson and Guigó 2014). To search for potential TE insertions, we analyzed the SChLAP1 Iso. 1 sequence using RepeatMasker (http://www.repeatmasker.org/), focusing on SChLAP1 Iso. 1 hereafter as it is the most abundant isoform of SChLAP1 (Prensner et al. 2013). We identified two long terminal repeats (LTRs) within SChLAP1: LTR12C (DFAM accession DF0000402), within exon 1 (nucleotides 1-322; 95.3% of exon 1) and THE1B (DFAM accession DF0000818) within exon 7 (nucleotides 1123-1436; 46.9% of exon 7) (Fig. 1B, Fig. S1)(Storer et al. 2021). LTRs occur at the 5′— and 3′—ends of retroviral genomes and are involved in viral processes such as replication. After integration into the host genome, the central components of the retroviral genome may be removed across evolution by recombination, such that only the LTR sequences remain (so-called solo-LTRs) (Babaian and Mager 2016; Johnson 2019). LTRs constitute over 8% of the human genome, and these sequences may be epigenetically silenced or, in contrast, “exapted” (i.e., co-opted) by the host genome for various functionalities including as promoters/enhancers (Babaian and Mager 2016; Johnson 2019). The occurrence of these sequences in SChLAP1 suggests that some of SChLAP1 function may be the result of such exaptation from these insertions and resultant RNA sequence/structure.

Our BLAST search results are consistent with evidence of retroviral insertion in exons 1 and 7. LTR12C, derived from human endogenous retrovirus 9 (HERV-9), is a Hominoidea (i.e., ape)-specific insertion. Our BLAST search only observed this sequence in *Homo sapiens*, *Gorilla gorilla gorilla*, *Pan troglodytes*, *Pan paniscus, Nomascus leucogenys*, and *Hylobates moloch* genomes, all of which are of Hominoidea phylogeny (Schoch et al. 2020; Storer et al. 2021). In comparison, the BLAST results did not identify LTR12C insertions nor the SChLAP1 exon 1 sequence within the *Colobus angolensis palliates*, *Cercocebus atys*, or *Mandrillus leucophaeus* genomes (Fig. 1A); these results are consistent with known phylogeny as these species are Old World monkeys and thus of non-hominoidea phylogeny (Schoch et al. 2020). The other retroviral insertion in SChLAP1 (THE1B) belongs to the MaLR (Mammalian Apparent LTR-Retrotransposons) family of repeat elements. In contrast to LTR12C, THE1B is specific to Simiiformes (i.e., simians), which contains both Hominoidea and Old-World monkeys (Schoch et al. 2020; Storer et al. 2021). In turn, a complete exon 7 sequence was observed in both lineages in our BLAST search. Our results indicate that THE1B resulted in the formation of approximately half of exon 7. Given that both LTR12C and THE1B insertions are selectively found in Hominoidea and Simiiformes, respectively, we believe these data may explain why SChLAP1 homologs were not observed in lower order species, e.g., mice, in our BLAST search.

Using the USCS Genome Browser (Kent et al. 2002), we found that the LTR12C sequence alignment to the 5′—end of human SChLAP1 not only overlaps with the majority of SChLAP1 exon 1 but also includes the promoter region of the *SChLAP1* gene (Fig. S2). Potential retroviral incorporation within the *SChLAP1* promoter has been mentioned in previous work (Prensner et al. 2013; Babaian and Mager 2016), and this analysis suggests the insertion may have also introduced novel sequence/structures into the 5′—end of the SChLAP1 transcript.

### Sequence and bioinformatic analyses identify potentially functional SChLAP1 substructures

To complement our phylogenetic analysis, we used additional sequence and bioinformatic analyses to identify regions of SChLAP1 that may be crucial to its function. To begin, we used ScanFold 2 (Andrews et al. 2022) to predict thermodynamically favorable and likely functional RNA secondary structures within SChLAP1. Specifically, ScanFold 2 folds a given transcript in sliding folding windows, shuffles the sequence of each window (100 times each in our case), and thereafter calculates the folding energy of each shuffled sequence. The folding energies for the native versus shuffled sequences are compared via z-score (defined as the average minimum free energy (MFE) of the shuffled sequences subtracted from the MFE and divided by the standard deviation), where z-scores of −1 and −2 indicate a folding window 1 and 2 standard deviations more stable than the shuffled folds, respectively. Previous work has found that regions with significantly lower energies of folding in the native versus shuffled sequence are often enriched in functional roles (Andrews et al. 2018; Andrews et al. 2022). From this work, we identified multiple SChLAP1 structures with significantly low z-scores (Fig. 2, top), including: 1) predicted stem loops in 5′—end of SChLAP1 (i.e., containing the LTR12C insertion); 2) the junction between exon 2 and exon 5 of SChLAP1 (hereafter referred to as the E2-E5 junction); 3) a predicted helix from nucleotides 989-1056 (within exon 7), and; 4) an adjacent predicted three-way junction (3WJ, nucleotides 1141-1213) and stem-loop (nucleotides 1224-1242), both located within exon 7 and the THE1B insertion. These analyses support the presence of potentially functional RNA structures within SChLAP1 on the basis of their favorable sequence-based folding properties. The above RepeatMasker analysis also suggests that some of these structures may have been derived through retroviral insertion.

**Figure 2.**
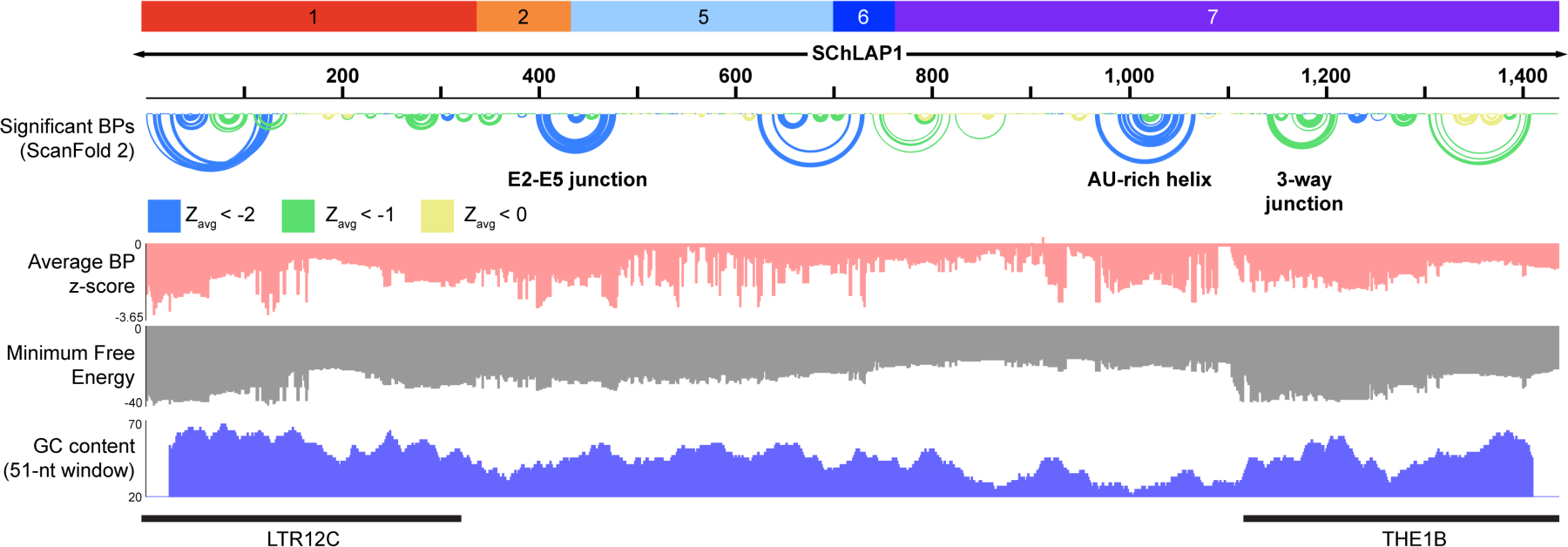
ScanFold 2 (Andrews et al. 2022) prediction of SChLAP1 Isoform 1. Top: Significant base-pairs identified using ScanFold 2. Second and third row depicted average base-pair z-score and minimum free energy of folded windows calculated by ScanFold. Bottom row shows GC content calculated over 51 nucleotide windows. Exon coordinates of SChLAP1 Iso. 1 are depicted along with coordinates of LTR12C and THE1B insertions. Figure generated in Integrative Genomics Viewer (Robinson et al. 2011).

Additionally, we calculated the GC content of SChLAP1 in 51-nucleotide sliding windows to identify regions of structure correlated with the more stable hydrogen bonding provided by canonical GC base pairs versus AU base pairs (Fig. 2, bottom). Overall, SChLAP1 comprises 44.5% GC, but contains windows as high as approximately 70% GC. Several ScanFold-predicted structures occur in such high-GC regions; however, we observed that one highly predicted structure (nucleotides 989-1056) occurs within a remarkably GC-depleted region of the transcript. We refer to this predicted structure as the AU-rich helix where it appears in our analysis below.

### Development of a secondary structure model for SChLAP1 *in vitro* using SHAPE-MaP

Our work so far has indicated that there are thermodynamically predicted functional structures within the SChLAP1 RNA transcript, some of which were derived from retroelements inserted into the primate lineage. We next set out to generate an experimentally informed secondary structure model of SChLAP1 using SHAPE-MaP. We first began by generating an *in vitro* structure SChLAP1 model as a preliminary reference to evaluate the propensity of these structures to form both *in vitro* and in cells (analyzed below). To this end, we performed SHAPE-MaP and DMS-MaP on *in vitro* transcribed SChLAP1 Iso 1. Full-length SChLAP1 Iso. 1 RNA was isolated using a semi-native purification protocol, where both heat denaturing and harsh buffer exchanges are avoided, as previous work found this protocol superior in maintaining a homogenously folded RNA as compared to denaturing protocols (Somarowthu et al. 2015; Adams et al. 2019). We chose the SHAPE reagent 5-nitroisatoic anhydride (5NIA) for its enhanced cell permeability compared to other SHAPE reagents and precedence for use of 5NIA in prostate cancer cells (Busan et al. 2019), which enabled comparable use of 5NIA in both *in vitro* and *in cellulo* experiments.

The *in vitro* secondary structure model of SChLAP1 was generated with SHAPE and DMS probing, which were performed with independent batches (i.e., transcribed, purified, and probed on different days) of SChLAP1 RNA. As SChLAP1 is >1,400 nucleotides, which hinders sequencing analysis on Illumina®-based platforms, we generated four overlapping amplicons (positions 1-500, 400-903, 800-1300, and 1200-1436) from the probed RNA to facilitate sequencing. These four amplicons were independently reverse transcribed (SuperScript II and Mn^2+^ for SHAPE probing, TGIRT for DMS) and amplified for sequencing. The SHAPE sequencing data was processed using the SHAPEMapper (Busan and Weeks 2018) and SuperFold (Smola et al. 2015b) pipelines, whereas the DMS-MaPseq program was used to process the DMS data (Zubradt et al. 2017) (see Methods). In both SHAPE and DMS experiments, we observed a lack of chemical reactivity in a poly(A) stretch of SChLAP1 (position 1088-1104), which is consistent with work from Kladwang et al. showing that chemical modifications in poly(A) regions are bypassed by reverse transcriptases and result in incongruous chemical modification frequency and mutational profiling results (Kladwang et al. 2020). To avoid biasing our resultant model as predicted by SuperFold/RNAStructure, we manually set this poly(A) stretch to “undefined” in the input .shape and .map files for all SHAPE-informed structure predictions of SChLAP1; setting these nucleotides to undefined removes all chemical reactivity data from these nucleotides such that only RNA folding dynamics are used to predict base-pairing, thereby excluding these data from biasing the predicted secondary structure.

The MFE model (Fig. S3A) and predicted base-pairs (Fig S4A) of *in vitro* SChLAP1 reveal a complex structural landscape, including significantly base-paired regions of varying secondary structure types as well as large single-stranded regions. Pseudoknot prediction was performed as published (Smola et al. 2016; Wan et al. 2022) but were not incorporated into the structure models reported herein due to lack of evidence for pseudoknot formation in SChLAP1 in our downstream *in cellulo* analysis (see Supplemental Text 1 and Fig. S5). The DMS data showed good agreement with the *in vitro* SHAPE data (Fig. S3A) and resulted in greater modification of adenosines and cytidines over guanosines and uridines as expected (Fig. S3B). We also observed agreement for several local structures between our SHAPE-informed partition functions and ScanFold models (Fig. S4A). These include the aforementioned AU-rich helix (nucleotides 989-1056), the 3WJ (nucleotides 1141-1213) and adjacent stem-loop (nucleotides 1224-1242), and the E2-E5 junction (nucleotides 398-478). The agreement between these models gives independent support for the formation of these various substructures in *in vitro* SChLAP1 as predicted by ScanFold 2.

### *Ex cellulo* SHAPE probing of SChLAP1 reveals cell-derived RNA structures

As our *in vitro* chemical probing data supported the presence of functionally important structures within the SChLAP1 RNA, we next set out to generate a secondary structure model for cell-derived SChLAP1 RNA to identify whether these structures or others fold in a physiologically relevant context. We performed SHAPE probing of endogenous SChLAP1 isolated from LNCaP prostate cancer cells, a common model cell line with high SChLAP1 expression as compared to normal prostate cells and cell lines (Prensner et al. 2013) and with a known metastatic phenotype. Due to the retroviral insertion in exon 7, which overlapped completely with amplicon 4 used in our *in vitro* probing, we extended the coordinates of amplicon 4 from what was used in our *in vitro* experiments to avoid amplification of other THE1B-containing transcripts (see Methods).

We began our in-cell structure modeling with RNA that was gently isolated from LNCaP cells and Proteinase K treated (i.e. *ex cellulo* RNA) using previously established protocols to extract cellular RNA without heat-denaturing (Smola et al. 2015a; Smola et al. 2016). Two independent replicates of *ex cellulo* probing of SChLAP1 yielded reactivity profiles with good agreement (Pearson’s R = 0.72). The resultant secondary structure arc diagrams (Fig. 3A and Fig. S4B,C) and MFE model (Fig. 4, see below) revealed a complex structure for cell-derived SChLAP1, with an array of predicted structures across the length of the transcript. We also observed agreement among several structures identified in our *ex cellulo* SHAPE data, *in vitro* SHAPE data, and ScanFold data (Fig. S4), including the E2-E5 junction, AU-rich helix, and 3WJ, giving independent support for the formation of these structures and their ability for form in cells.

**Figure 3.**
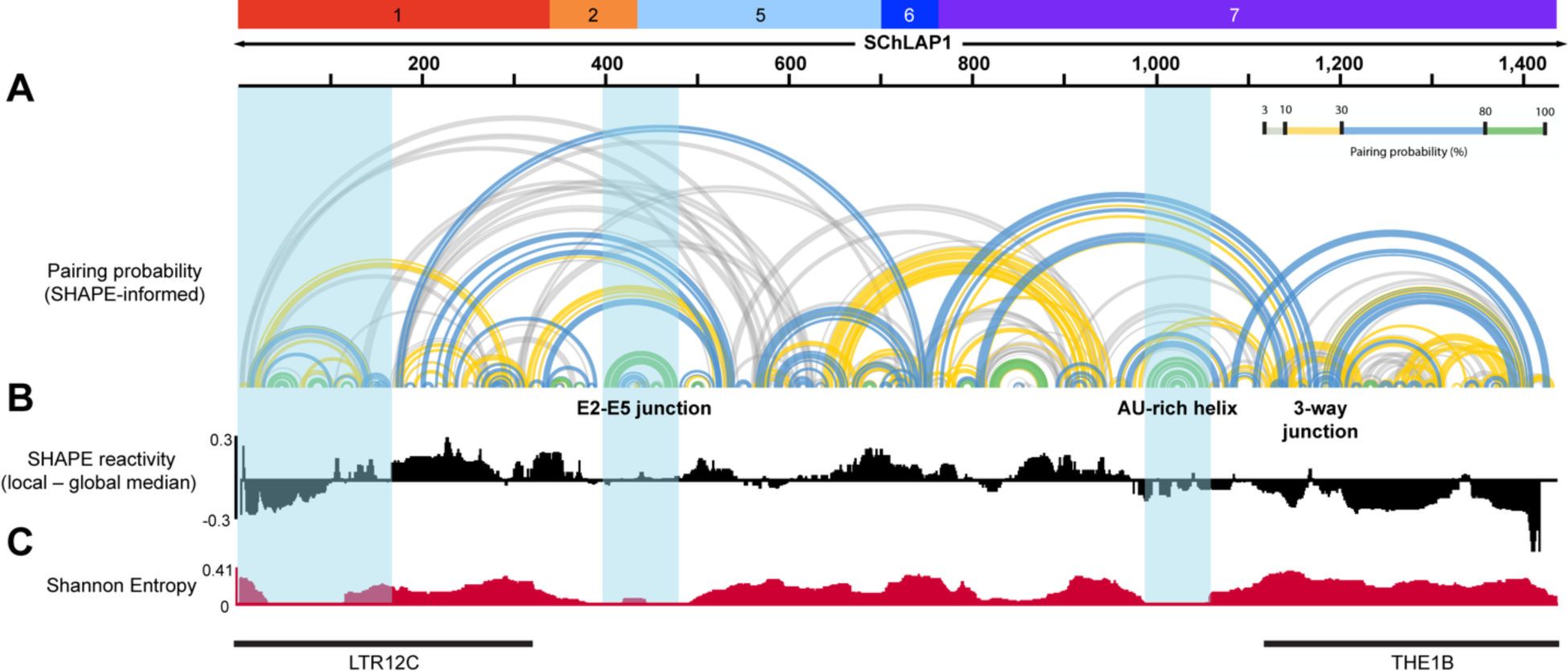
SChLAP1 secondary structure generated through *ex cellulo* SHAPE-MaP from LNCaP cells (representative of two independent replicates). A) Arc diagram showing the SHAPE-informed base pair predictions across *ex cellulo* SChLAP1. Arc diagram of second replicate shown in Fig. S4C. B) Median SHAPE reactivity of 51-nucleotide windows across the length of SChLAP1 subtracted from the global median (SHAPE = 0.31). C) Shannon entropy plot across SChLAP1. Blue boxes denote low SHAPE/Shannon regions that were reproducible across both biological replicates. Figure generated in Integrative Genomics Viewer (Robinson et al. 2011).

**Figure 4.**
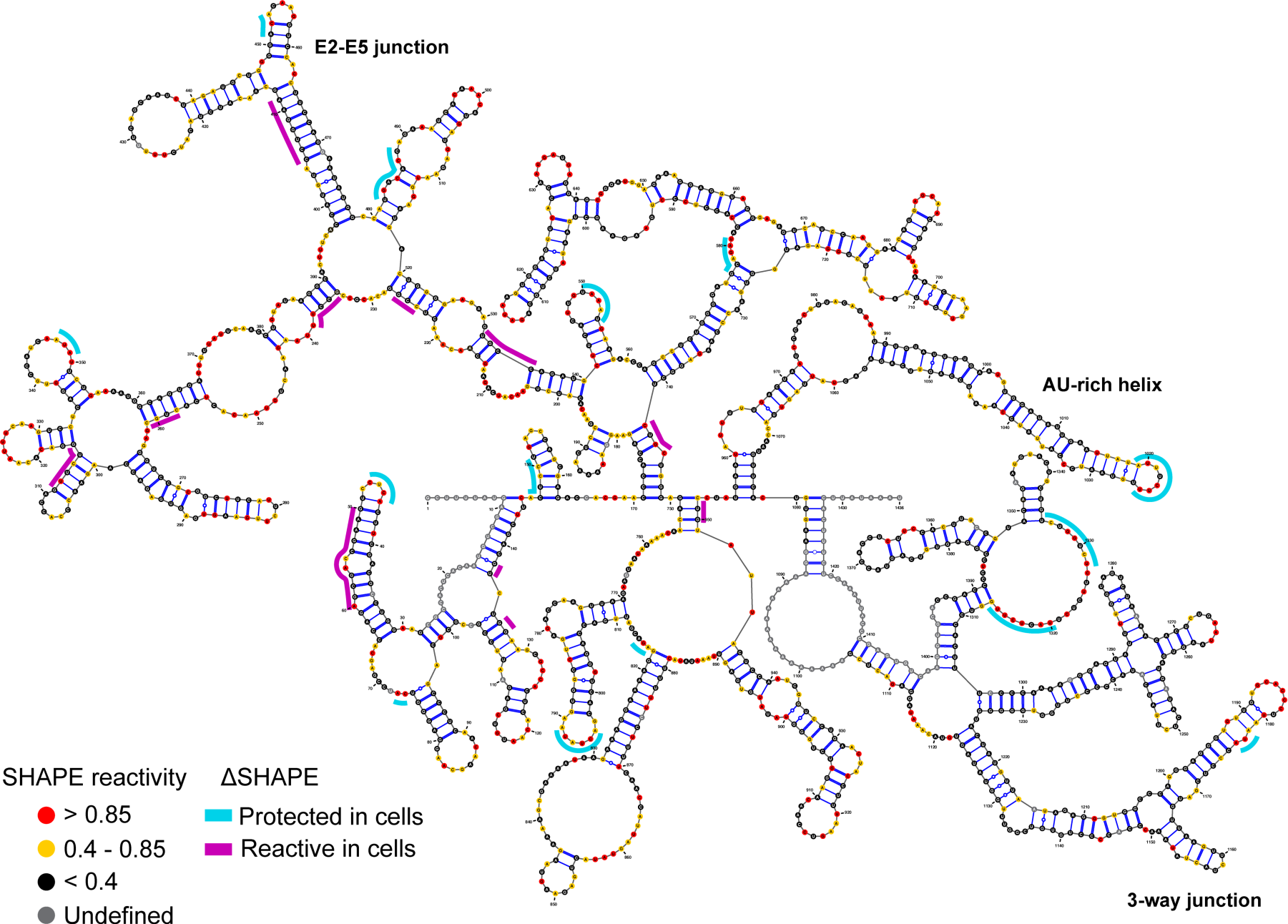
ΔSHAPE analysis of SChLAP1. Significant in-cell protected or enhanced regions identified in both replicates of ΔSHAPE (see Methods) are mapped over a representative MFE structure of *ex cellulo* SChLAP1. SChLAP1 structure visualized in VARNA (Darty et al. 2009).

We further analyzed our *ex cellulo* SHAPE data to identify putative functional structures within cell-derived SChLAP1. Work by Siegfried et al. characterizing HIV-1 genomic RNA found that RNA structural elements classified as 1) highly structured (i.e., low SHAPE reactivity) and 2) well-determined (i.e., low Shannon entropy), which are referred to as low SHAPE/Shannon (lowSS) regions, were significantly enriched in previously unknown functional roles (Siegfried et al. 2014). This metric has since been used to identify novel functional regions, including in the Dengue virus RNA genome (Dethoff et al. 2018), XIST lncRNA (Smola et al. 2016), and HCV genome (Wan et al. 2022). To identify if SChLAP1 contained lowSS regions, we calculated the SHAPE reactivity (51 nucleotide local median compared to global median, Fig. 3B) and Shannon entropy (51 nucleotide windows, Fig. 3C) to identify lowSS sites. In our SChLAP1 model, we identified three lowSS regions: nucleotides 1-165 (within LTR12C insertion), nucleotides 398-478 (E2-E5 junction), and nucleotides 989-1056 (AU-rich helix) (Fig. 3, blue bars). These lowSS regions agree well with ScanFold predicted structures (Fig. 2).

Additionally, we observed that the nucleotides between the abovementioned lowSS regions (i.e., nucleotides 166-397, which contains portions of exon 1 and 2, and nucleotides 479-988, which contains portions of exons 5 and 7 and the entirety of exon 6) display both elevated SHAPE reactivities and Shannon entropies relative to the median (Fig. 3B and 3C). This result indicates that these regions are generally single-stranded and/or conformationally heterogenous. Consistent with these results, the arc diagrams of the two *ex cellulo* probing replicates vary considerably in these regions (Fig. S4C). In contrast, downstream from the lowSS site located at nucleotides 989-1056 (i.e., nucleotides 1057-1436), we observe low SHAPE reactivity alongside high Shannon entropy, indicating a highly structured but poorly predicted and/or conformationally heterogenous structure. This highly structured region correlates well to the THE1B insertion within SChLAP1 (nucleotides 1123-1436).

We next compared the consistency of our *in vitro* and *ex cellulo* SHAPE profiles using Pearson correlation coefficients. We calculated the correlations using 51-nucleotide sliding windows for both replicates of our *ex cellulo* SChLAP1 against our *in vitro* SHAPE data. We observed regions of high correlation between our *ex cellulo* and *in vitro* data alongside regions of low/modest correlation (Fig. S6). In particular, SChLAP1 nucleotides 1-204, 318-477, and 960-1105 were the largest regions with high correlation between our *in vitro* and *ex cellulo* data (defined as having Pearson R ≥ 0.5 in both replicates, a threshold chosen heuristically based on previous comparisons of *in vitro* and *in/ex vivo* chemical probing studies (Frank et al. 2020; Sherpa et al. 2018; Manfredonia et al. 2020)). Thus, these regions are more likely to recapitulate a similar structure *in vitro* and *ex cellulo.* These regions also align with the aforementioned lowSS regions in our *ex cellulo* data (Fig. 3). We also identified smaller regions that show correlation between the *in vitro* and *ex cellulo* models, particularly within nucleotides 803-857, which corresponds to a predicted stem and 5′—end portion of a large, single-stranded loop in our *in vitro* (Fig. S3A) and *ex cellulo* (Fig. 4) MFE models.

### Identification of protein binding regions via *in cellulo* SHAPE

SChLAP1 has been documented to interact with several proteins (see Introduction). As our *in vitro* and *ex cellulo* work identified potentially functional secondary structures within SChLAP1 (in particular, ScanFold and lowSS regions) as well as single stranded regions that could additionally function in protein binding, we next performed *in cellulo* SHAPE probing to identify whether these structures persist in cells and/or function as putative protein binding regions that may be involved in any of the aforementioned interactions. For consistency with the *ex cellulo* experiments, *in cellulo* SHAPE probing was performed in the LNCaP cell line. The resultant SHAPE reactivity profiles between two independent replicates showed good agreement (Pearson’s R = 0.59). For reference, replicate *in cellulo* SHAPE probing experiments of endogenously expressed Xist lncRNA yielded a Spearman correlation coefficient of 0.50 (Smola et al. 2016). We then used the ΔSHAPE algorithm to identify protein binding regions/structures of SChLAP1 by comparing our *in cellulo* reactivity data and *ex cellulo* data, as the ΔSHAPE algorithm has previously identified protein binding sites in other lncRNAs and viral RNAs (Smola et al. 2016; Frank et al. 2020; Jones et al. 2020; Schmidt et al. 2020; Wan et al. 2022; Przanowska et al. 2022).

The results of two replicates of ΔSHAPE analysis are shown on a representative MFE structure from our *ex cellulo* probing (Fig. 4); each replicate (see Methods) is individually shown in Fig. S7. Across both replicates of ΔSHAPE analysis, we observed indications for protein binding (in-cell protections) in both structured regions and relatively unstructured regions as previously identified in our *ex cellulo* analysis. For example, the three lowSS regions in our *ex cellulo* model, i.e., the beginning of exon 1/LTR12C insertion, the E2-E5 junction, and the AU-rich helix, each contain *in cellulo* protected nucleotides, indicating that these structures likely participate in intermolecular interactions and suggesting a role for these highly predicted RNA structures in protein binding. Additionally, we observed significant in-cell protections within the remainder of exon 5, which was found to be relatively high in SHAPE reactivity and Shannon entropy and suggestive of an unstructured, single stranded region (Fig. 3). These data suggested that this region functions as a single stranded “landing pad” for protein binding, with similar regions observed in other RNAs such as Xist (Smola et al. 2016; Weeks 2021). Lastly, we observed significant in-cell protections in the low SHAPE, high Shannon 3′—end of SChLAP1 (within THE1B insertion), indicating that this region is also playing important roles for protein binding. In particular, these protections were observed in the aforementioned 3WJ (nucleotides 1141-1213, supported by ScanFold (Fig. 2) as well as our *in vitro*/*ex cellulo* predicted base-pairs (Fig. 3 and Fig. S4)) and a CU-rich single-stranded region (nucleotides 1313-1335, observed in our *in vitro*/*ex cellulo* SHAPE-informed MFE models (Fig. S3A, Fig. 4). As this region corresponds to the THE1B LTR insertion (Fig. 1B, Fig S1), these results suggest that the formation of this SChLAP1:protein recognition interface occurred *de novo* through a retroviral insertion within primate evolution.

Gratifyingly, several putative protein binding regions identified herein are consistent with previous work from other groups, supporting the validity of our ΔSHAPE data. For example, in-cell protections observed in exon 2 are consistent with recent work from Ji et al., where binding between SChLAP1 exon 2 and HNRNPL facilitated activation of the NF-κB pathway in glioblastoma (Ji et al. 2019). While exon 2 lacks the canonical poly-CA binding site for HNRNPL, which was previously observed for its RNA interactome in prostate cancer (Fei et al. 2017), a CA-rich region (nucleotides 341-349) overlaps with an in-cell protected region in our ΔSHAPE data (nucleotides 347-349) and may support weak or indirect interaction with HNRNPL. In addition, protein binding with exon 7 is in line with previous work from the Chinnaiyan group. Preliminary work from Sahu et al. revealed that a deletion of 250 nucleotides (from position 1001-1250) of SChLAP1 Iso. 1 inhibited SChLAP1-driven invasion and binding to the SWI/SNF complex (Sahu 2015). These coordinates overlap with two regions of in-cell protection, the former within the AU-rich helix and the latter within the predicted 3WJ in the THE1B insertion. Both sites are also contained within favorable ScanFold predicted structures (Fig. 2). While interaction with the SWI/SNF complex is contested (Raab et al. 2019), the role of this particular region in protein recognition is supported by our work.

### Analysis of unfolded regions in cellular SChLAP1 compared to *ex cellulo*

Alongside the above in-cell protections, we also observed regions of in-cell enhancement of reactivity (i.e., nucleotides with more reactive *in cellulo* as compared to *ex cellullo* data) by ΔSHAPE, indicating the unfolding of some structures in cells as compared to the *ex cellulo* model. In particular, in-cell enhancements appeared to be enriched at the 5′—end of the transcript and more limited in the remaining sequence (Fig. 4 and Fig. S7B). Within the first 500 nucleotides of SChLAP1 (i.e., the first 34.8% of sequence), 76 of 130 (58.5%) and 82 of 120 (68.3%) instances of in-cell enhancement are observed in replicates 1 and 2 of ΔSHAPE analysis, respectively (Fig. S7B). While the ΔSHAPE algorithm emphasizes robust local changes in SHAPE reactivity (Smola et al. 2015a; Smola et al. 2016), we hypothesized that the abundance of *in cellulo* enhancements called by ΔSHAPE at the 5′—end of SChLAP1 may be reflective of larger-scale structural changes. To analyze broader conformational changes between *ex cellulo* and *in cellulo* SChLAP1, we calculated the overall SHAPE reactivity change in 51-nucleotide windows between our *ex cellulo* and *in cellulo* samples as was employed with the Xist lncRNA (Smola et al. 2016). Several regions of significant SHAPE reactivity change were identified (Fig. S7C). Of particular note, we observed significant enhancement of SHAPE reactivity of exon 1 in cells, which was consistent with the multiple ΔSHAPE-identified *in cellulo* reactivity enhancements we observed (Fig. 4 and Fig. S7B). These results suggest that this region is largely unfolded in a cellular environment. Notably, these results contradict the ScanFold prediction (Fig. 2, top), high GC content (Fig. 2, bottom), *in vitro* probing data (Figs. S3A and S4A), and *ex cellulo* probing data (Figs. 3 and 4), where significant structure was indicated. Additionally, we identified several in-cell protections by ΔSHAPE in this region (Fig. 4 and Fig. S7B), indicating that, despite the larger-scale unfolding of this region, local sites of *in cellulo* protection occur and suggest participation in intermolecular interactions.

Similar to exon 1, we observed that the E2-E5 junction shows enhancement of SHAPE reactivity *in cellulo* by ΔSHAPE (Fig 4, Fig. S7B). However, our ΔSHAPE data indicate that the majority of in-cell enhancements in this region are located on the 5′—end of the E2-E5 junction. Despite these in-cell enhancements, the structure of the E2-E5 junction is still strongly predicted in the arc diagram for our in-cell data (Figs. S7A and S8A). We examined the SHAPE reactivities of the largest stems in this region (nucleotides 398-412 and 464-478, the former containing multiple in-cell enhancements), and observed statistically significant enhancement of SHAPE reactivity *in cellulo* compared to *ex cellulo* for the 5′—stem, but not the 3′—stem in this region (Fig. S8B). In addition, we observed ΔSHAPE *in cellulo* protections in a small stem-loop in the E2-E5 junction (nucleotides 449-461, Fig. 4), suggesting that protein binding occurs within this structure despite the presence of *in cellulo* enhancements elsewhere. From these data, we hypothesize that protein binding within the E2-E5 junction occurs primarily on the 3′—portion of the structure but not the 5′—portion. As a result, the 5′—stem of the structure, while putatively base-paired to the 3′—stem *in vitro* and *ex cellulo*, is more accessible to chemical modification *in cellulo*. We note that this proposed disruption would be distinct from the broader unfolding observed for exon 1/LTR12C insertion.

While the 5′—end of SChLAP1 appears to be generally unfolded in cells, we observed few ΔSHAPE *in cellulo* enhancements at the 3′—end of SChLAP1 (Fig. 4 and Fig. S7B). This result is supported by the presence of highly predicted base-pairs at the 3′—end rather than the 5′— end of SChLAP1 *in cellulo* (Fig. S7A). Thus, we believe that most of the structures at the 3′—end of SChLAP1, such as the AU-rich helix and 3WJ, remain folded in cells, allowing these RNA structures to function within cells as well-folded protein binding substructures.

### *In vitro* folding analysis of SChLAP1 3′**—**end reveals conformational heterogeneity in the THE1B LTR insertion

While our above work characterized SChLAP1 secondary structures and protein binding interfaces, we next set out to evaluate if any of these recognition interfaces participated in tertiary/higher order structural complexity. We specifically focused on the 3′—end of SChLAP1 for several reasons. First, this region showed generally low SHAPE reactivity, yet high Shannon entropy, indicating a structured yet poorly predicted region that may be due to structural complexity or heterogeneity. Consistent with these results, the correlation analysis between our *ex cellulo* and *in vitro* replicates (Fig. S6) showed differences in the correlation coefficients at the 3′—end of SChLAP1, indicating variability between experiments in this region. Given that ΔSHAPE analysis supports protein binding at the 3′—end of SChLAP1, we were interested in examining this region’s functionality as a structured protein binding region. Second, evidence of retroviral insertion in this region (i.e., THE1B) indicates that the RNA structures in this region may have emerged from an external source; indeed, recent evidence points to the role of transposable elements in forming *de novo* RNA structural domains. In particular, Feschotte and coworkers compared the predicted structures of 100 lncRNAs with high TE content (over 96% of sequence) to 100 low-TE content lncRNAs (less than 3% of sequence) and observed a greater propensity for statistically significant secondary structures in high-TE lncRNA as evaluated by a sequence randomization approach similar to ScanFold (Kapusta et al. 2013). While THE1B insertions have been previously analyzed for their roles In gene promoters and enhancers for both coding and non-coding genes (Lamprecht et al. 2010; Young et al. 2013; Dunn-Fletcher et al. 2018), they have not to our knowledge been analyzed in terms of RNA structure. Thus, this evidence for a structured RNA domain corresponding to the THE1B insertion, along with evidence that the THE1B insertion remains folded in cells while the LTR12C insertion is unfolded in cells, led us to hypothesize that this insertion could have resulted in the formation of a complex structural landscape that may be additionally related to its protein binding capability.

To assess the folding landscape of the 3′—end of SChLAP1, specifically within exon 7 and the THE1B insertion, we used native gel electrophoresis. This method was chosen due to its flexibility to accommodate RNA of various sizes and ability to assess varying conditions (e.g., salt identity and concentration) in an efficient manner (Woodson and Koculi 2009). Using our *ex cellulo* model along with additional structure predictions in RNAstructure (Bellaousov et al. 2013) as guidance for substructures to study, two strongly predicted structures from the 3′—end of SChLAP1 were chosen for analysis by native gel electrophoresis: Arm A (nucleotides 949-1116), which contains the above-mentioned AU-rich helix, and Arm B (nucleotides 1117-1428), which contains the 3WJ and corresponds to the THE1B insertion (Fig. S9A).

We began by looking at two variables, magnesium (Mg^2+^) concentration and annealing method, and their impact on Arm A/B migration by gel electrophoresis. Mg^2+^ is a divalent cation known to be crucial for the stabilization of RNA tertiary structures (Butcher and Pyle 2011) whereas the method of RNA annealing, specifically whether the RNA is snap-cooled or slow-cooled, may modify the resultant conformational landscape of the RNA population. In our initial examination of these variables, wherein Arm A/B were snap- or slow-cooled in filtration buffer and thereafter incubated with varying Mg^2+^ concentrations (protocol designed for consistency with our chemical probing above), Arm A migrated at a consistent distance and as one conformer via native gel regardless of either variable (Fig. 5, top). This result indicates that neither the annealing protocol nor the Mg^2+^ concentration significantly impacts the resultant structure of Arm A. We note that this finding does not rule out more subtle conformational changes that may occur within Arm A. In contrast, for Arm B we observed an unexpected trend between Mg^2+^ concentration and annealing protocol (Fig. 5, bottom). When Arm B is snap-cooled, Mg^2+^ addition significantly alters the conformational space of Arm B, favoring the formation of two central conformations over a lower and higher conformation observed at low magnesium concentrations. However, when Arm B is slow-cooled, this Mg^2+^ dependence is greatly reduced, and Arm B migrates in one major conformer throughout the magnesium titration.

**Figure 5.**
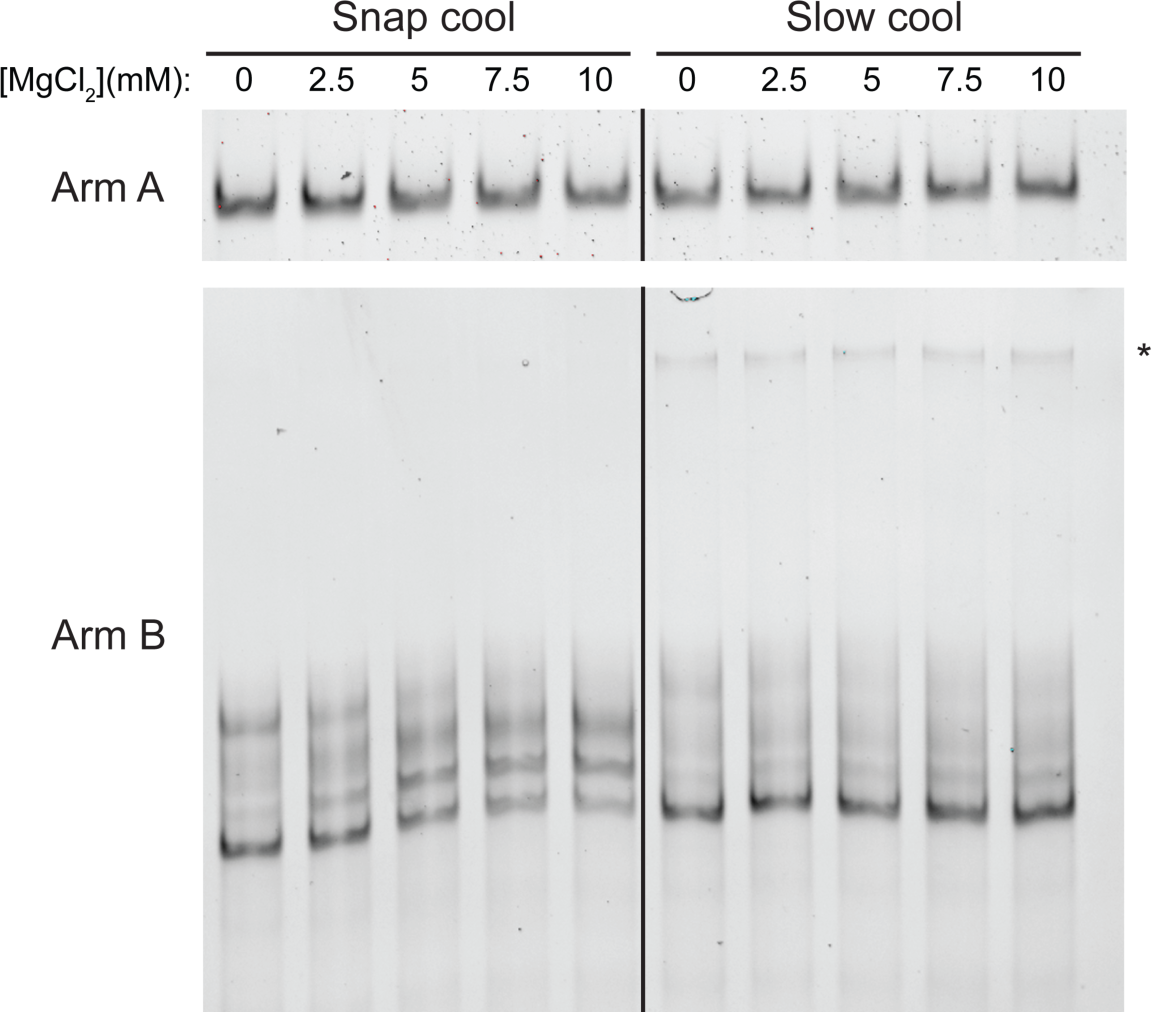
Native gel electrophoresis of Arm A and Arm B with varying annealing protocols and magnesium concentrations. RNA was snap or slow cooled in filtration buffer (50 mM K-HEPES pH 7.5, 150 mM KCl, 0.1 mM EDTA) before incubation with the given Mg^2+^ concentration. Asterisk denotes putative Arm B dimer favored in slow cooling. RNA was purified using the “kit-purified” approach. Arm A gel representative of two biological replicates, and Arm B gel is representative of three biological replicates.

Next, we evaluated how snap- and slow-cooling impact the migration of Arm B when incubated with divalent cations besides Mg^2+^. We evaluated the migration of Arm B in response to 5 mM Ba^2+^, Sr^2+^, Ca^2+^, Mn^2+^, and Ni^2+^ (all as metal chlorides). We chose these divalent metals as their roles in RNA folding and stability have been assessed in previous studies, particularly for RNA G-quadruplexes and the *Tetrahymena* ribozyme, and they cover an array of chemical properties including radius, charge density, and hydration enthalpy (Koculi et al. 2007; Balaratnam and Basu 2015). As above, Arm B was snap- or slow-cooled in filtration buffer before incubation with the indicated divalent metal. For snap-cooled Arm B, we found that some metals, such as Ba^2+^ and Sr^2+^, resulted in Arm B migrating primarily in the lowest conformer (i.e., fastest moving band) observed with 0 Mg^2+^. However, when incubated with other metals, such as Mn^2+^ and Ni^2+^, the additional, higher apparent conformations of Arm B are preferred (Fig. 6A). Again, this trend was not observed for slow-cooled Arm B (Fig. 6B), where the lowest conformer is observed regardless of metal incubation.

**Figure 6.**
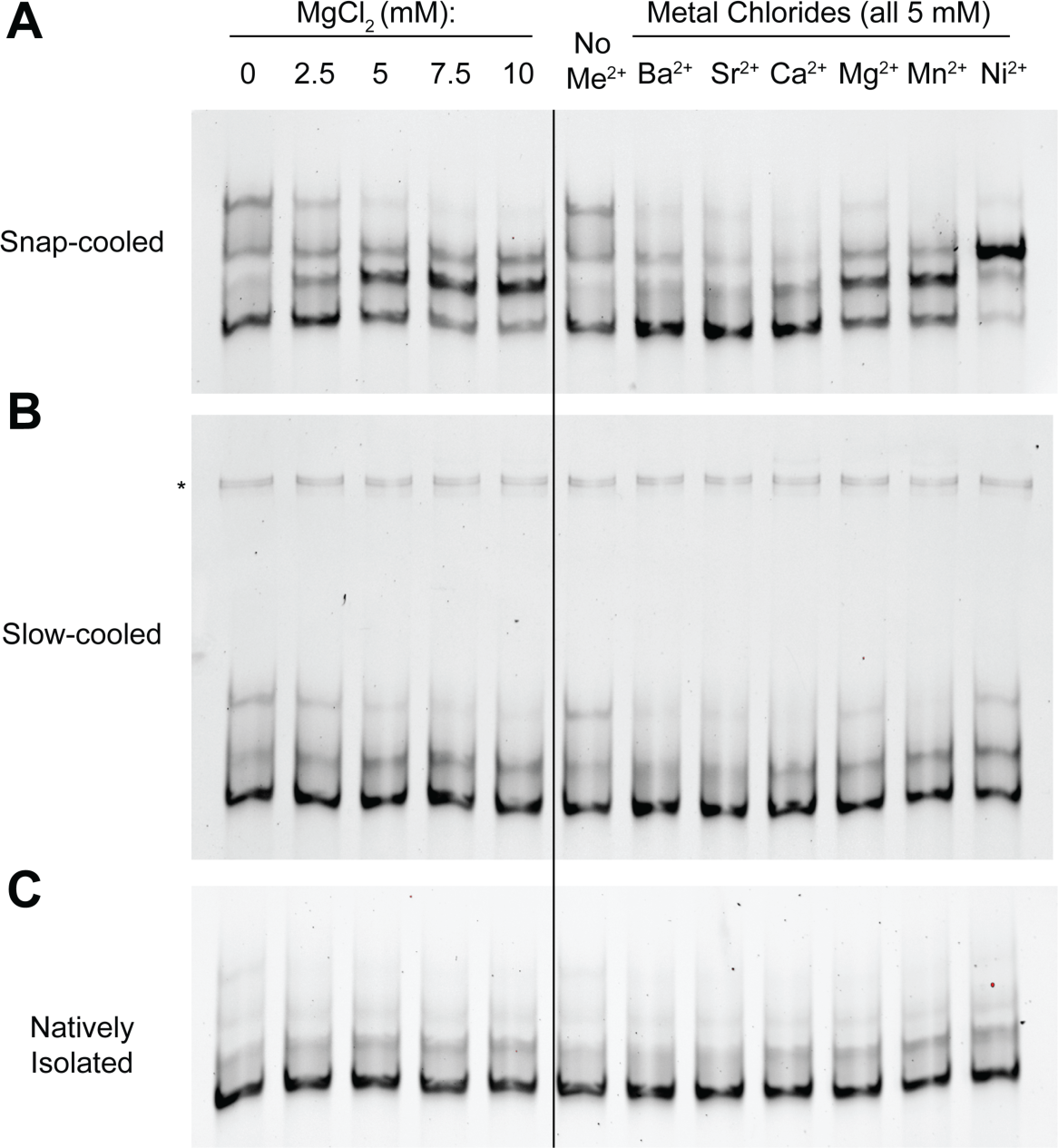
Native gel electrophoresis of Arm B with various divalent metals and annealing conditions. Arm B was prepared by snap-cooling (A), slow cooling (B), or native isolation (C) in filtration buffer (see Methods) before incubation with divalent metals. Asterisk denotes dimer favored during slow cooling. Black line through gels is for reference only; magnesium titration and incubation for other divalent metals was run on the same gel for each panel. RNA was generated using FPLC purification, and each gel was run in biological duplicate (kit-purified of snap- and slow-cooled Arm B run in triplicate, Supplemental Fig. 9B).

Given that changes in Arm B migration in the presence of divalent metal were dependent on snap- or slow-cooling, we next compared natively purified Arm B, where the co-transcriptionally folded structure is preserved. We isolated Arm B using the same native purification approach that we used for full-length SChLAP1 and thereafter incubated the RNA with the aforementioned divalent metals. We observed that the metals did not induce a significant change in migration for natively folded Arm B (Fig. 6C), i.e. Arm B migrated as one major conformer regardless of metal incubation. We additionally confirmed that the differences inherent to the native purification workflow did not impact the resultant RNA conformational landscape by snap- and slow-cooling Arm B that had previously been natively purified. The same trends were observed for these samples by native gel: snap-cooling Arm B that was previously natively purified resulted in the abovementioned metal-dependence that was not observed if natively purified Arm B was slow-cooled (Fig. 6 and Fig. S9B).

Together, our native gel data suggest that metal recognition by Arm B is significantly altered by annealing method, wherein snap-cooled Arm B shows altered migration by divalent metals depending on their identity, but slow-cooled and natively folded Arm B show largely consistent migration regardless of divalent metal. These results indicate that 1) a structural difference within Arm B occurs between snap- and slow-cooling 2) slow-cooling rather than snap-cooling much more effectively mimics the native fold of Arm B, and 3) that divalent metals alter the conformational landscape of a non-native Arm B fold. These data give support for structural complexity within this region of SChLAP1 *in vitro*, which may in turn explain the elevated Shannon entropy observed in our *ex cellulo* SHAPE data (Fig 3C). As stated above, this complex structure additionally appears to function as a protein binding region.

### THE1B insertion within SChLAP1 may form a G-quadruplex, but is not sufficient to explain its conformational heterogeneity *in vitro*

Mg^2+^ is generally thought to favor RNA compaction, but in our experiments, an increase in apparent size is observed with increasing Mg^2+^. The order of divalent cation identity in increasing the apparent size of snap-cooled Arm B is reminiscent of recent work of divalent metal recognition of G-quadruplexes, where metals such as Ni^2+^ induced an unfolded conformation of a G-quadruplex at lower concentrations over other metals such as Ba^2+^ (Balaratnam and Basu 2015). To evaluate potential G-quadruplex formation, we snap- and slow-cooled Arm B in a modified filtration buffer wherein the concentration of the G-quadruplex stabilizing cation K^+^ (in the form of KCl) was varied from 0 to 150 mM prior to separation by native gel. We observed a significant change in migration for both annealing conditions with increasing K^+^, indicating the presence of a K^+^-dependent structure regardless of annealing protocol (Fig. 7A). As a control, we studied the impact of K^+^ on Arm A migration, and we observed no change in migration in response to K^+^ for Arm A (Fig. S10A). This result suggests that a G-quadruplex forms within Arm B in both annealing protocols and that G-quadruplex formation may be a component of the conformational heterogeneity in Arm B/THE1B.

**Figure 7.**
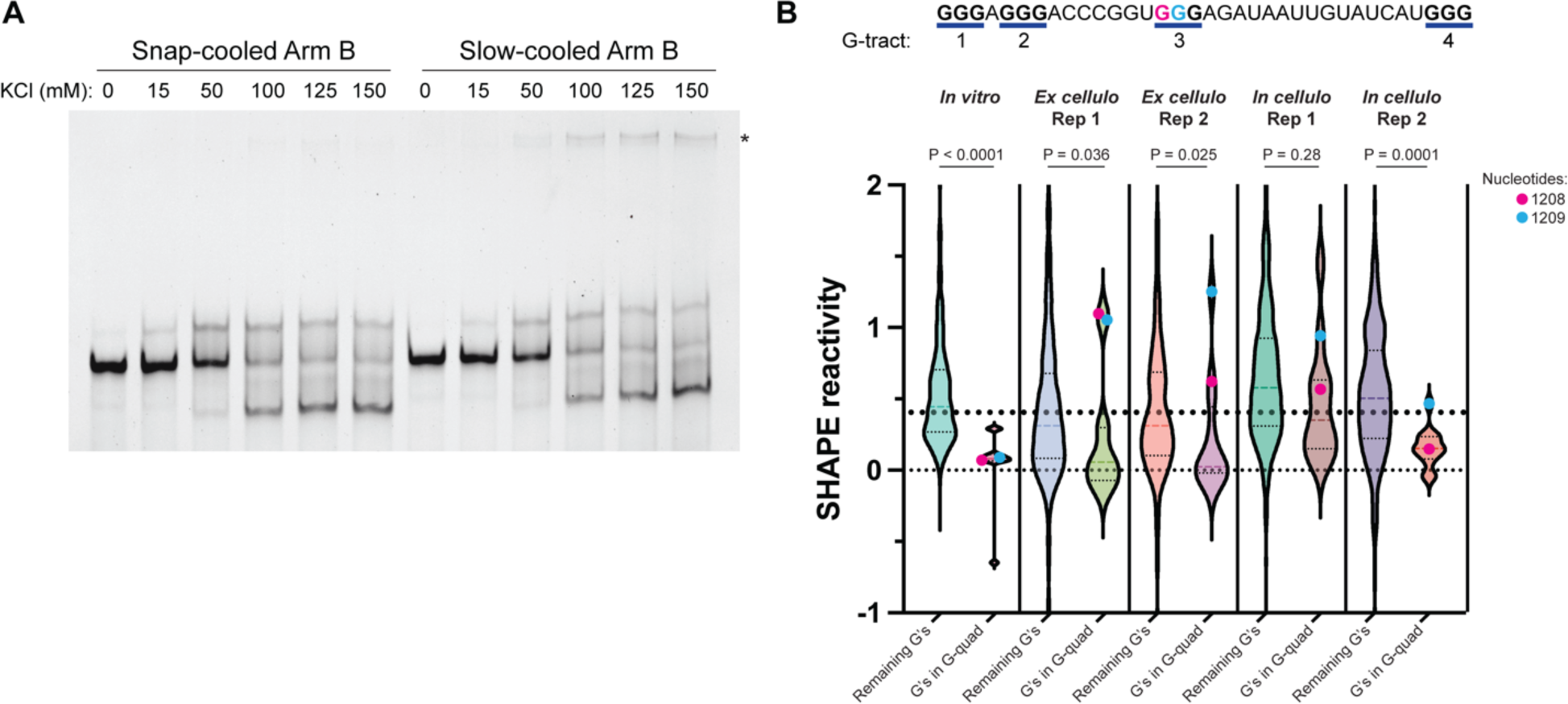
Analysis of G-quadruplex formation of Arm B by native gel electrophoresis and SHAPE. A) Native gel electrophoresis of kit-purified Arm B in response to varying KCl concentrations and annealing conditions. Gel is representative of two biological replicates. B) Top: sequence of predicted G-quadruplex in Arm B by QGRS Mapper (Kikin et al. 2006). G-tracts are numbered, and guanosines predicted to participate in G-quadruplex are in bold. Bottom: violin plots comparing the reactivities of guanosines in predicted G-quadruplex to the remaining guanosines in each SHAPE probing experiment reported herein. Reactivities below −1 or above 2 were not plotted here but were included in statistics. A line at SHAPE = 0.4 is shown for reference as SHAPE reactivities above this value are unfavorable for base-pairing (Hajdin et al. 2013). Significance was calculated using a Kolmogorov-Smirnov test for each experiment. Nucleotides 1208 and 1209 are indicated by color coding.

To further investigate potential G-quadruplex formation, we used QGRS Mapper (Kikin et al. 2006) against the SChLAP1 Iso. 1 sequence to search for potential G-quadruplex forming regions. Searching with the default parameters, which allows G-quadruplex formation within a 30-nucleotide window, we identified four lowly predicted potential G-quadruplex forming regions (Fig. S10B, left). However, extending the maximum allowed G-quadruplex to 45 nucleotides facilitated the prediction of a 3-tract G-quadruplex (tract referring to the number of guanosines in a row per strand of the putative G-quadruplex) located within Arm B (nucleotides 1194-1228) (Fig. S10B, right). This predicted G-quadruplex was the only predicted G-quadruplex with three G-tracts, which we hypothesized could confer added stability to this fold compared to the other predicted G-quadruplexes in SChLAP1, which only contained two G-tracts. In addition, this predicted G-quadruplex would have two large loops, one of which (nucleotides 1211-1225) was predicted by RNA structure (Bellaousov et al. 2013) to form a stem-loop structure itself (Fig. S10C).

This computational result supports our K^+^-dependent native gel studies, where evidence of G-quadruplex formation was observed in Arm B (and thus the THE1B insertion). However, we note that this G-quadruplex overlaps with the coordinates of the ScanFold/SHAPE-informed 3WJ (nucleotides 1141-1213) and adjacent stem-loop (nucleotides 1224-1242), indicating that a non-quadruplex conformation is also thermodynamically predicted.

### Analysis of SHAPE data indicates a structured but non-quadruplex conformation of Arm B/THE1B insertion in cell-derived SChLAP1

Given the above indication of a G-quadruplex structure in Arm B of SChLAP1 *in* vitro, and its overlap with favorably predicted non-quadruplex secondary structures, we evaluated our *in vitro and in/ex cellulo* SHAPE data for evidence of G-quadruplex formation. Recent work from Schneekloth and coworkers using the probe 2-methylnicotinic acid imidazolide found that guanosines participating in a G-quadruplex within the *NRAS* mRNA 5′ UTR were generally low in SHAPE reactivity *in vitro*, supporting their structuredness (Balaratnam et al. 2023). We analyzed the SHAPE reactivity of the predicted G-quadruplex within Arm B identified above (nucleotides 1194-1228) to evaluate its structuredness and indications of quadruplex formation. In four of our five probing experiments, we observe a statistically significant decrease in SHAPE reactivity of the guanosines participating in the putative G-quadruplex compared to the SHAPE reactivity of all remaining guanosines (Fig. 7B), suggesting that the guanosines in this potential G-quadruplex are indeed functioning in an RNA structure that hinders SHAPE reactivity. While statistical significance was not obtained for in-cell replicate 1, the data trends in the same direction and may be consistent with heterogeneity of this structure in *in cellulo* RNA as indicated above. However, we observed that two residues in this predicted G-quadruplex (nucleotides 1208 and 1209, within G-tract 3) showed a large enhancement in SHAPE reactivity in our *ex cellulo* data as compared to our *in vitro* data (Fig. 7B). While our *in vitro* data showed low SHAPE reactivity in each G-tract, consistent with our K^+^-dependent native gels for Arm B that indicated G-quadruplex formation (Fig. 7A), the heightened SHAPE reactivity in our *ex cellulo* data for G-tract 3 in cell-derived SChLAP1 suggests non-quadruplex structure. This trend is also observed in our *in cellulo* data, although with greater variation between the replicates (Fig. 7B).

A non-quadruplex structure for this region is supported by our representative *ex cellulo* MFE structure (Fig. 4), where a 3WJ-type structure is supported by ScanFold (Fig. 2). In this model, the abovementioned nucleotides 1208 and 1209 are base-paired across a 1-nucleotide bulge, which may introduce more flexibility in these nucleotides despite their base-pairing and thus yield heightened SHAPE reactivity for these nucleotides in cell-derived RNA. Thus, while the data suggest that Arm B of SChLAP1, which encompasses the THE1B insertion, has a propensity to form G-quadruplexes, especially *in vitro*, this conformation appears less likely to form in cell-derived SChLAP1. Nonetheless, the combined data supports the SHAPE reactivity predictions of a well-structured but conformationally heterogeneous region the 3′—end of SChLAP1.

## DISCUSSION

The study of lncRNA structure-function relationships is of great interest to the scientific community from both a basic science perspective as well as therapeutic one. While three-dimensional analysis of lncRNA structure by methods such as X-ray diffraction (XRD), nuclear magnetic resonance (NMR), and cryo-electron microscopy (cryo-EM) are limited, in large part due to lncRNA size and conformational dynamics, chemical probing methods such as SHAPE-MaP combined with phylogenetic analyses have provided insight into these large biomolecules, exemplified by XIST (Lu et al. 2016; Smola et al. 2016; Jones et al. 2019). Importantly, these chemical probing methods have often informed and/or reproduced three-dimensional structure findings and have even expanded upon them as seen in rRNA (Mustoe et al. 2019) and Dengue virus genomic RNA (Dethoff et al. 2018).

The phylogenetic and structural analyses presented herein have produced the first targeted secondary structure model of the lncRNA SChLAP1, which serves as a foundation for further biochemical and biophysical analyses. First, sequence-based phylogenetic analysis of SChLAP1 human and NHP sequences revealed a high degree of sequence conservation, particularly among exons 1, 2, 5, and 7 (Fig. 1A), suggesting function in these regions. Complementary to our phylogenetic work, RepeatMasker analysis revealed the insertion of two LTRs within SChLAP1 (Fig. 1B and Fig. S1): LTR12C and THE1B. While outside the scope of this work, we hypothesize that reactivation of a retroviral promoter, derived from LTR12C, may be responsible for SChLAP1 overexpression in prostate cancer. Consistent with this idea, recent work from Feng and coworkers uncovered reduced methylation of the SChLAP1 promoter in castration-resistant prostate cancer samples over benign prostate samples (Zhao et al. 2020).

Examining SChLAP1 *in cellulo* revealed significant disruption of structures at the 5′—end of the transcript relative to other data, particularly within the LTR12C insertion (Fig. S7C). This result indicates that these regions are unfolded in the cellular environment despite having significant ability to form RNA structures as indicated by ScanFold (Fig. 2) and *in vitro*/*ex cellulo* probing (Fig. S3, Figs. 3 and 4), which is supported by work from Rouskin and coworkers that reported active unfolding of RNA structures *in vivo* (Rouskin et al. 2013). As ΔSHAPE simultaneously identified in-cell protections at the 5′—end, it is likely that regional intermolecular contacts (e.g., RNA-protein interactions) co-occur with this observed unfolding. Whether this unfolded conformation of the LTR12C insertion relates to SChLAP1’s ability to participate in intermolecular interactions in cells is worthy of further investigation. In contrast, while we observe disruption of the E2-E5 junction in cells (Fig. S8), we hypothesize that this is the result of protein binding occurring primarily on the E2-E5 3′—end, thereby freeing the E2-E5 5′—end for SHAPE modification. We expect that additional studies into SChLAP1, particularly proteomics, will provide more insight into the interactome of the substructures predicted here, including the E2-E5 junction, AU-rich helix, and 3WJ.

The putative protein recognition sites identified in this work are in agreement with previous studies that have characterized SChLAP1-protein interactions, namely with HNRNPL in glioblastoma (Ji et al. 2019) and SWI/SNF in prostate cancer (Prensner et al. 2013; Sahu 2015). For the former, HNRNPL has been implicated in prostate cancer progression as promoting cell growth and alternative splicing; HNRNPL knockdown significantly diminished LNCaP cell growth, but did not impact RWPE-1 cell growth, indicating cancer specific functions for HNRNPL (Fei et al. 2017). However, we note that SChLAP1 binding by HNRNPL was not observed in the HNRNPL RIP-seq data reported in this study (Fei et al. 2017), potentially suggesting an indirect or weak interaction.

With regard to SWI/SNF, our results align with work by Sahu and co-workers (Sahu 2015), where the SChLAP1 Iso. 1 1001-1250 nucleotide region was necessary to bind the SWI/SNF complex subunit SNF5; critically, these nucleotides exhibited several in-cell protections in our data and overlap with the AU-rich helix and 3WJ in our structure model (Fig. 4). Additionally, other works have uncovered functional roles for RNA:SWI/SNF recognition, including X-inactivation and nuclear paraspeckle formation amongst others (Cajigas 2015; Kawaguchi et al. 2015; Wang et al. 2015; Hu et al. 2016; Lino Cardenas et al. 2018; Huang et al. 2019; Jégu et al. 2019; Grossi et al. 2020) despite known promiscuous binding of SWI/SNF across the genome and transcriptome (Cajigas et al. 2015; Raab et al. 2019; Grossi et al. 2020; Skalska et al. 2021). Additional mechanistic characterization of SChLAP1:SWI/SNF interaction is needed to understand the specific role of SChLAP1:SWI/SNF in prostate cancer progression.

The 3′—end of SChLAP1 appears to remain folded in cells, as evidenced by lack of in-cell enhancements (Fig. 4 and Fig. S7). Further, this region also showed indications for protein binding by ΔSHAPE and includes a significant portion of the aforementioned 1001-1250 region found critical for SWI/SNF binding (Sahu 2015), indicating that this region may be functionally relevant for prostate cancer progression. Despite these data, which indicate significant structure and function, the 3′—end of SChLAP1 is also high in Shannon entropy and therefore has a poorly predicted or heterogenous structure (Fig. 3C). To further analyze this conformational heterogeneity, we used native gel electrophoresis to characterize this region, named Arm B, derived from the THE1B insertion. We observed a complex structure fold within Arm B as evidenced by 1) changes in migration in the presence of some divalent metals, but not others, 2) the dependence of annealing protocol for these changes in migration, and 3) the role of potassium in altering Arm B migration. We observed no evidence of these trends in our control structure, Arm A, indicating that these trends are specific to Arm B. This is the first work to our knowledge that has analyzed the RNA conformational landscape of a THE1B insertion, which is notable given that THE1B insertions are common in the human genome (over 18,000 copies) (Storer et al. 2021). These results indicate that the insertion of this retroviral domain may have introduced a structured yet conformationally diverse protein binding hub within SChLAP1 that may facilitate a role in promoting aggressive prostate cancer. These results also suggest that THE1B insertions may participate in the function of other RNAs where the insertion is transcribed.

While the landscape within Arm B may be in part due to G-quadruplex formation *in vitro*, our analysis suggests that other, non-quadruplex containing conformations are accessed preferentially in cell-derived RNA. We expect that higher resolution methods, such as chemical crosslinking or cryo-EM, will better define the contacts within Arm B that determine this conformational landscape. In addition, we expect that further investigation of the Arm B structure, including probing with new methods and reagents *in vitro* and *in cellulo*, as well as obtaining 3D structure information, will also give greater insight into the contacts that are changed between its native and nonnative conformations, as was recently employed for the *Tetrahymena* ribozyme (Li et al. 2022).

In conclusion, we observed that SChLAP1 has a complex secondary structure, containing a multitude of intramolecular interactions both *in vitro* and *in cellulo*, many of which are potentially functional sites of RNA:protein recognition that may facilitate its role in prostate cancer progression. We believe the putative structure-function relationships identified here will facilitate both fundamental understanding of prostate cancer progression and the development of specific therapeutic strategies against SChLAP1 for prostate cancer treatment. In particular, our work identifies single-stranded regions of SChLAP1 that may be amenable to antisense oligonucleotide (ASO)/siRNA-based therapies alongside more structured regions, which we hypothesize may be amenable to small molecule-based therapies (Warner et al. 2018; Costales et al. 2020; Falese et al. 2021). We expect that future work will capitalize on the potential for SChLAP1 as a diagnostic marker and therapeutic target to ultimately yield better patient outcomes for those with aggressive prostate cancer.

## MATERIAL AND METHODS

### Cell culture

LNCaP cells were obtained from the Duke University Cell Culture Facility and were authenticated with short tandem repeat and mycoplasma testing. Cells were grown in RPMI 1640 media (Gibco) with 10% fetal bovine serum (FBS, Gibco) at 37°C in a humidified atmosphere with 5% CO_2_. All experiments were performed on cells less than 20 passages before retrieving a fresh vial from cryopreservation.

### Phylogeny Analysis

BLAST search (Zhang et al. 2000) was performed using isoform 4 of SChLAP1 as it contains all possible exons of human SChLAP1. The following search paramaters were used to putative identify primate homologs of SChLAP1: Database = Reference RNA Sequences (refseq_rna) (O’Leary et al. 2016), Optimize for: more dissimilar sequences (discontiguous megablast). Figure 1 was generated using megablast comparing each identified primate sequence (Table S1) to isoform 4.

### RepeatMasker analysis of SChLAP1

RepeatMasker (http://www.repeatmasker.org/) was performed on the SChLAP1 Isoform 1 sequence with the following settings: Search Engine: rmblast; Speed/Sensitivity: default; DNA Source: Human. No lineage annotation options were selected. Alignment options: no alignments returned; masking options: repetitive sequences replaced by strings of N’s; contamination check: no contamination check; repeat options: masked interspersed and simple repeats; artifact check: report *E. coli* insertion elements artifacts; Matrix: RepeatMasker choice.

### Structure Analysis with ScanFold

Identification of significant local RNA structures was performed with ScanFold 2 (Andrews et al. 2022) using the following parameters: window size: 120; step size: 1; randomizations: 100; temperature: 37; filter value: −2; global refold: false; competition: true.

### DNA Template and Primers

All oligos were purchased from Integrated DNA Technologies (IDT). For generation of full-length SChLAP1 isoform 1 for *in vitro* transcription, the SChLAP1 sequence was inserted downstream of a bacteriophage T7 RNA polymerase promoter and upstream of the BAMHI restriction site. Plasmid growth was performed by transformation into NEB5α competent cells (New England Biolabs, USA) following manufacturer’s instructions and subsequent selection on LB agar plates with ampicillin (100 µg/mL final) for overnight growth at 37°C. A single colony was propagated in LB broth with ampicillin selection, and plasmids were isolated using the Qiagen Plasmid Kit. The plasmid was linearized using BAMHI-HF (New England Biolabs, USA) following manufacturer protocol. Linearized plasmid was purified using Qiagen DNA Mini Kit. PCR reactions were performed using Q5 High-Fidelity DNA polymerase (New England Biolabs, USA) and purified using DNA Clean and Concentrator 5 kit (Zymo). To generate Arm A and Arm B, primers unique to the Arm A or Arm B were ordered, with the forward primer bearing a 5′ overhanging T7 promoter sequence such that the T7 promoter would be incorporated during PCR. See SI Table 2 for all construct sequences, SI table 3 for primer sequences, and SI Table 4 for RNA sequences.

### *In vitro* SHAPE probing of SChLAP1

*In vitro* transcription (IVT) for SChLAP1 Isoform 1 was completed following the procedure from Adams *et al*. with some modifications (Adams et al. 2019). T7 RNA polymerase was a generous gift from Blanton Tolbert’s lab (Case Western). No RNase inhibitor was used in any of the steps. IVT of these constructs was performed by mixing: 200 µL 10X Transcription buffer (400 mM Tris-HCl pH 8.0, 100 mM NaCl, 120 mM MgCl_2_, 20 mM spermidine, 0.1% Triton X-100), 200 µL rNTPs (25 mM equimolar mix), 25 µL T7 RNA polymerase (custom preparation), 25 µL Yeast Inorganic Pyrophosphatase (YIPP, 2 kU/mL, New England Biolabs, USA), 50 µg PCR-amplified DNA template; 100 µL molecular biology grade DMSO (5% final), and nuclease-free water up to 2 mL. This mixture was aliquoted into 1.5 mL Eppendorf tubes at 500 µL each and incubated at 37°C for 2-4 hours. DNase I, Proteinase K treatments, and RNA concentration were followed as outlined in Adams *et al* (Adams et al. 2019). Following concentration of the reaction with 100 kDa MWCO Amicon filter to a final volume of 1 mL, size exclusion chromatography was performed at room temperature using Bio-Rad NGC FPLC. A Cytiva (formerly GE Healthcare) HiPrep Sephacryl 16/60 S-500 column was used for SChLAP1 WT. An isocratic method was employed using 1X filtration buffer (FB; 50 mM K-HEPES, pH 7.5, 150 mM KCl, 100 μM EDTA pH 8.0). Prior to use, columns were washed with 3 column volumes (CV) of 1:1 RNase ZAP (Ambion) followed by 3 CV nuclease-free water (DEPC-treated MilliQ water), and finally equilibrated with 3 CV 1X FB. Flow rates were between 0.5–0.75 mL/min, and 0.5 mL fractions were collected. RNA peaks were monitored using UV_255_ absorbance. The largest absorbance of the product peak and two surrounding fractions were used for downstream experiments. Nanodrop and/or Qubit confirmed RNA concentration, and purity was verified using agarose gel electrophoresis before proceeding. SChLAP1 isoform 1 was typically purified by SEC at approximately ∼90 ng/µL for all probing reactions and were diluted with 1X FB if necessary to achieve this concentration.

RNA from the FPLC was maintained at room temperature prior to addition of MgCl_2_ to a final concentration of 5 mM final (i.e. 2.5 µL of 10X magnesium concentration in FB was added to 20 µL RNA for a total of 22.5 µL). These reactions were then incubated at 37°C for 30 minutes. 5 nitroisatoic anhydride (5NIA, Millipore Sigma) was prepared at in DMSO immediately before probing. Following incubation with MgCl_2_, 5NIA was added to each reaction, flicked to mix, and incubated at 37°C for 10 minutes. At 10 minutes, each reaction was quenched by adding 33% final volume BME (Sigma-Aldrich) and placed on an ice block pre-chilled to −20°C. Prior to ethanol precipitation, Sephadex G-50 columns (Cytvia/GE Healthcare) were used to remove the hydrolyzed 5NIA reagent following the manufacturer instructions. SChLAP1 isoform 1 was probed in one biological replicate with SHAPE reagent and an independent biological replicate with DMS.

### DMS Chemical Probing

*In vitro* DMS probing was completed following protocols from the Rouskin lab (Zubradt et al. 2017). PCR, IVT, and FPLC purification of SChLAP1 Iso. 1 was completed as described above. Following FPLC purification, RNA samples were adjusted to 300 mM HEPES and 5 mM MgCl_2_ before incubation at 37°C for 30 minutes prior to DMS probing. DMS (EMD Millipore) was diluted into 100% molecular biology grade ethanol for a final concentration of 2% upon addition to the RNA sample and was prepared immediately prior to use to limit oxidation. DMS was incubated with the RNA at 37°C for 5 minutes while shaking at 500 rpm. The reaction was quenched after 5 minutes by adding 33% final volume BME (Millipore Sigma) and placed on an ice block pre-chilled to −20°C.

### *In Cellulo* SHAPE

In-cell SHAPE probing was performed with the reagent 5NIA in the LNCaP cell line following previous protocols (Smola et al. 2015a; Smola et al. 2016; Busan et al. 2019). Specifically, LNCaP cells were plated at 5 x 10^5^ cells per well in a 6-well plate and grown for approximately 2 days. On the day of the experiment, cells were washed with 1 mL warm PBS (Geneclone), and 900 uL complete media was added to each well. To control wells, 100 µL anhydrous DMSO (Invitrogen) was added. For SHAPE-probed wells, 100 µL of freshly prepared 250 mM 5NIA was added. Gentle swirling was used to evenly distribute the SHAPE reagent. The reactions were incubated in an incubator at 37°C for 15 minutes. After probing, the media was removed, and cells were washed with 1 mL warm PBS. Total RNA from each reaction was extracted with TRIzol reagent and resuspended in nuclease-free water. The solutions were treated twice with DNAse I and thereafter purified using RNA Clean/Concentrator 25 columns (Zymo).

### *Ex Cellulo* SHAPE

*Ex cellulo* SHAPE was performed with 5NIA following previous protocols with RNA isolated from the LNCaP cell line (Smola et al. 2015a; Smola et al. 2016; Busan et al. 2019). Cells were grown to approximately 80% confluency in a T-75 flask. The cells were trypsinized, pelleted, and resuspended in ice-cold PBS. Four million cells were centrifuged and resuspended in 2.5 mL freshly prepared lysis buffer (40 mM Tris-HCl (pH 7.9), 25 mM NaCl, 6 mM MgCl_2_, 1 mM CaCl_2_, 256 mM sucrose, 0.5% (vol/vol) Triton X-100, 1,000 U/ml RNase inhibitor (Promega), and 450 U/mL DNase I (Roche)), and rocked at 4°C for 5 minutes. This mix was centrifuged at 2250 xg for 2 minutes at 4°C. The resultant pellet (nuclei) was resuspended in 2.5 mL Proteinase K buffer (40 mM Tris-HCl (pH 7.9), 200 mM NaCl, 1.5% w/v SDS, and 500 μg/ml Proteinase K (Millipore Sigma)) and treated for 45 minutes at approximately 20°C. After Proteinase K treatment, 2.5 mL phenol:chloroform:isoamyl alcohol (25:24:1) pre-equilibrated with 1.1X refolding buffer (55 mM K-HEPES pH 7.5, 165 mM KCl, 5.5 mM MgCl_2_, 0.11 mM EDTA; chosen for consistency with *in vitro* work) was added, and the sample was vortexed and centrifuged at 4000 xg for 15 minutes at 4°C. The aqueous phase was isolated, and the pre-equilibrated phenol:chloroform:isoamyl alcohol extraction was repeated. Thereafter, the aqueous phase was mixed with 2.5 mL chloroform, vortexed, and centrifuged as above. The aqueous phase was isolated, and the chloroform extraction was repeated. The resultant aqueous phase was buffer exchanged into 1.1X refolding buffer using PD-10 columns (Cytvia) following the gravity protocol. The eluate was separated into two tubes of equal volume (approximately 1.7 mL) and incubated at 37°C for 30 minutes. DMSO was added to the control tube, and 5NIA (25 mM final) was added to the SHAPE-treated well. Reactions were incubated at 37°C for 15 minutes. Control and probed RNA was isolated by ethanol precipitation and resuspended in 87 µL nuclease-free water. DNAse treatment and column clean-up were performed identically to the *in cellulo* samples.

### RNA Reverse Transcription

To sequence SChLAP1 for mutational profiling, we utilized an amplicon workflow wherein SChLAP1 was sequenced in four overlapping amplicons of maximum 500 nucleotides (approximately 600 upon addition of sequencing adaptors; amplicon sequences provided in SI Table 2). Each amplicon overlapped with the adjacent amplicons via a 100 base-pair window. Thus, four unique reverse transcription reactions (one for each amplicon) were performed for each condition (mock or SHAPE probed) to generate sequencing profiles for the full SChLAP1 sequence for each experiment. For generating sequencing profiles for cell-derived amplicon 4, SChLAP1 was initially reverse transcribed with the amplicon 4 reverse primer followed by PCR with the amplicon 4 reverse primer and the amplicon 3 forward primer. The subsequent product was gel purified and underwent PCR amplification with step 1 primers specific to the original amplicon 4 coordinates. Sequences for all primers are provided in SI Table 3. Reverse transcription reactions were performed with blunt-end primers (i.e., no overhangs).

Reverse transcription of probed RNA was adapted from previous protocols (Adams et al 2019; Smola et al. 2015b). For *in cellulo* probed RNA, approximately 2000 ng total RNA (mock or 5NIA probed) was mixed with 1 µL 10 µM reverse primer (unique to each amplicon) and brought to 10 µL final in nuclease-free water. For *ex cellulo* probed RNA, 750 ng RNA was used as the input for RT. Primers were annealed by incubating at 65°C, followed by incubation on ice for 15 minutes. To these mixes, 8 µL 2.5X MaP buffer was added (125 mM Tris-HCl pH 8, 187.5 mM KCl, 25 mM DTT, 1.25 mM dNTP mix, 15 mM MnCl_2_), followed by 2 uL SuperScript II (Invitrogen). Reactions were incubated at 25°C for 10 minutes, 42°C for 3 hours, 70°C for 15 minutes, and then stored at −20°C. Reverse transcription reactions were cleaned-up using DNA Clean and Concentrator 5 columns (Zymo) before proceeding to PCR and library preparation.

For reverse transcription of *in vitro* probed SChLAP1 WT, 1 µg of RNA was diluted into 16 µL of nuclease-free water and divided such that 250 ng RNA went into each RT reaction. To each reaction, 1 µL of 1 µM respective primer was added and incubated at 65°C for 5 minutes before the reaction was placed on ice. Then, 8 µL of 2.5X MaP Buffer was added and incubated at 42°C for 2 minutes before addition of 1 µL SuperScript II Reverse Transcriptase (ThermoFisher) and incubation at 42°C for 3 hours. Samples were heat inactivated at 70°C for 15 minutes.

Reverse transcription for *in* vitro DMS-treated samples was adapted from previous protocols (Zubradt et al. 2017), 0.5 μg RNA was mixed with 2 μL 10X FSB (500 mM Tris pH 8.0, 750 mM KCl, 100 mM DTT), 1 μL dNTPs (10 mM equimolar mix), 1 μL TGIRT-III Reverse Transcriptase (200 U/μL, InGex), 1 μL of 10 μM respective reverse primer, 1 μL of 1M DTT, and brought to 20 uL final with nuclease-free water. The reactions were incubated at 65°C for 90 minutes and inactivated at 85°C for 5 minutes.

### SHAPE library preparation

Following reverse transcription, SChLAP1 amplicons were initially amplified using blunt end primers (SI Table 3). Thereafter, library preparation was performed using a two-step PCR reaction to add on Illumina primers following the amplicon workflow outlined previously (Smola et al. 2015b). Step 1 PCR primer sequences are provided in SI Table 3. PCR was performed using the Q5 high-fidelity polymerase (New England Biolabs). After each PCR step, amplicons were gel purified as needed using 1% or 2% agarose EX E-gels (Invitrogen) and Zymoclean Gel DNA Recovery Kit (Zymo) and thereafter quantified using a Qubit dsDNA High Sensitivity (HS) assay kit (Invitrogen). After completion of library preparation, samples were submitted to the Duke University School of Medicine Sequencing and Genomic Technologies Shared Resource. Samples were pooled and sequenced on an Illumina MiSeq Sequencer using Reagent Kit v3 (2 x 300 bp).

DMS library preparation.

Following reverse transcription (outlined above), DMS-probed samples were amplified into dsDNA using blunt-end primers. These samples were submitted to the Whitehead Institute for Biomedical Research Genome Technology Core for tagmentation prior to sequencing as described above.

### Bioinformatics Pipeline

SHAPE reactivity profiles, error estimates, mutation counts, and sequencing depths were obtained using the ShapeMapper pipeline (v 2.1.5) developed by the Weeks lab (UNC-Chapel Hill) (Busan and Weeks 2018). All default parameters were used. Samples were filtered at ≥1,000 nucleotide read depth.

After running ShapeMapper for each amplicon, amplicons within the same experiment were normalized using the box-plot method (Low and Weeks 2010). Mutation rates and error values from the output reactivity profiles were manually averaged within the overlapping regions for each experiment. The 5 nucleotides upstream of the reverse primer binding sites were not considered for averaging due to low fidelity of SuperScript II at reverse transcription start sites, consistent with previous work (Przanowska et al. 2022). If a nucleotide had a defined mutation rate in the overlapping windows for one amplicon, but not the other (e.g., defined in amplicon 3, but not in amplicon 4), the mutation rate from the amplicon where it was defined was taken forward without averaging. Thereafter, the normalization scale was calculated. Specifically, the interquartile range (IQR) of the averaged data was calculated, and nucleotides with mutation rates 1.5X the IQR were removed from calculation of the normalization scale. The normalization scale was calculated as the average of the top 10% of the remaining nucleotides. The mutation rates and standard errors of all nucleotides were divided by this normalization scale to give a normalized .map file. The 5′ primer binding site, 3′ primer binding site (with 5 nucleotides upstream this site excluded), and any nucleotides that were undefined in starting map files were set as undefined in these updated map files. In addition, a poly(A) stretch (nucleotides 1088-1104) was set to −999 due to bypass of chemical probing data in poly(A) stretches (Kladwang et al. 2020).

As it was previously established that 5NIA has a bias for reactivity with adenosines, the map files generated above were rescaled using scale_nuc.py from the Weeks group using the command --reagent 5NIA (Busan et al. 2019). After re-scaling, these final map files were used for downstream analyses. These reactivities, along with DMS reactivities (below) are provided in Supplementary Data I.

For DMS probing, mutation counts were calculated in the DMS-MaPSeq program from the Rouskin Group (Zubradt et al. 2017). DMSO-only mutation counts were subtracted from DMS-probed mutation counts. The mutation counts for adenosine and cytosine nucleotides were normalized following the above protocol. Normalized reactivities were manually overlaid onto the SHAPE-informed minimum free energy structure of *in vitro* SChLAP1 Iso. 1 (Fig. S3A).

### Structure modeling

Arc diagrams and Shannon entropies for full-length SChLAP1 were generated using SuperFold (v1.0)(Smola et al. 2015b). Minimum free energy structure models of full-length SChLAP1 (*in vitro* and *ex cellulo*) were generating using the RNAstructure command line with SHAPE data as input and the command --maxdistance 600 used (Reuter and Mathews 2010). For smaller constructs (e.g. Arm A and Arm B), the RNAstructure webserver was used with default parameters (Bellaousov et al 2013).

Pseudoknot predictions was carried out using ShapeKnots in the RNAstructure command line with default parameters (Hajdin et al. 2013). SHAPE reactivities from *ex cellulo* and *in vitro* data were divided into 600 nucleotide windows separated by 100 nucleotides. Pseudoknots predicted in multiple folding windows were considered (Smola et al. 2016; Wan et al. 2022). SHAPE reactivities were manually inspected for compatibility with pseudoknot formation and additionally checked against *in cellulo* reactivity (see Supplemental Text 1).

G-quadruplex prediction was performed using QGRS Mapper (Kikin et al. 2006), with default parameters used except for as mentioned in the Results section for the maximum length parameter.

### Identification of protein binding sites by ΔSHAPE

Map files from the *ex cellulo* and *in cellulo* experiments were analyzed using ΔSHAPE following default parameters (Smola et al. 2015). Given the known sensitivity of ΔSHAPE to hyperreactive nucleotides (Smola et al. 2016; Schmidt et al. 2020), we developed a protocol to exclude such hyperreactive nucleotides by identifying and removing the top and bottom 0.1% of SHAPE reactivities from each experiment (corresponding to the top and bottom 2 nucleotides for a given .map file). We note that these nucleotides were maintained for structure modeling and identification of lowSS regions. The 95^th^ percentile of the reactivities for each input map file did not differ by more than 3%, so no additional rescaling was performed for ΔSHAPE analysis as has been done previously (Smola et al. 2016; Schmidt et al. 2020). As previously performed for Xist (Smola et al. 2016) each replicate was handled independently; that is, replicate 1 of *in cellulo* probing was compared to replicate 1 of *ex cellulo* probing, and replicate 2 of *in cellulo* probing was compared to replicate 2 of *ex cellulo* probing. The most significant regions of in-cell enhancement or protection (depicted in Fig. 4) were identified as follows: nucleotides reproduced between both replicates of ΔSHAPE analysis were identified. From these nucleotides, neighboring ΔSHAPE-identified nucleotides from each replicate (tolerating gaps of 1 nucleotide) were merged together to generate a final coordinate of significant protection or enhancement (e.g. nucleotides 45, 46, and 47 in replicate 1 and 44, 45, and 46 in replicate 2 were merged to 44-47 for Fig. 4).

### Comparison of SHAPE reactivities for *in cellulo* vs *ex cellulo* SChLAP1

For comparison of larger reactivity changes between *ex cellulo* and *in cellulo* SChLAP1, SHAPE reactivities for *in cellulo* or *ex cellulo* SChLAP1 were averaged over 51-nucleotide windows, and these windowed values were subtracted (*in cellulo* – *ex cellulo*) as done previously (Smola et al. 2016). As with ΔSHAPE analysis, the top and bottom 0.1% of SHAPE reactivities were removed to facilitate comparison. Regions of significant reactivity were identified as 30 nucleotides or greater where the absolute value of the reactivity change was higher than the median of the absolute value of all reactivity changes, similar to previous work (Smola et al. 2016). Only regions that were supported in both replicates were considered significant.

### Identification of well-folded RNA structures

Calculation of regions with low SHAPE reactivity and low Shannon entropy was carried out as previously described (Smola et al. 2015b; Wan et al. 2022). Median SHAPE reactivities and Shannon entropies were calculated over 51-nucleotide sliding windows. Thereafter, the global medians of the SHAPE reactivities and Shannon Entropies were subtracted from those local values. Regions at least 40 nucleotides in length (with gaps of up to 6 nucleotides) where the local median SHAPE/Shannon values were lower than the global median SHAPE/Shannon were considered potential LowSS sites. Only LowSS sites reproducible between replicates (i.e. overlapping nucleotides, or replicates where nucleotides on opposite sides of a predicted stem were called) were considered in our final analysis. The coordinates of LowSS sites were expanded to include an entire partition-function predicted structure.

### Correlation Analyses and Statistics

Pearson correlation coefficients were calculated in Graphpad Prism (Version 9) or Microsoft Excel (Version 16). Before comparison, nucleotides within primer binding sites or undefined reactivities were removed. If a nucleotide had an undefined reactivity in one SHAPE profile, it was removed from both profiles in the comparison before calculation of correlation coefficients. For comparison between *in vitro* and *ex cellulo* SChLAP1 (Fig. S6), the outlier removal approach as used in ΔSHAPE was also employed. All t-test statistics were performed in Graphpad Prism.

### *In vitro* transcription of Arm A and Arm B

Arm A and Arm B were *in vitro* transcribed overnight at 37°C in the following conditions: 2.5 mM rNTP mix, 25 mM MgCl_2_, 40 mM Tris-HCl pH 8.0, 2.5 mM spermidine, 0.01% Triton X-100, 10 mM DTT, 0.5 U/mL Yeast inorganic pyrophosphatase (New England Biolabs), approximately 125 µg/mL T7 RNA polymerase, and either 5% or 10% DMSO for Arm A or Arm B, respectively. After *in vitro* transcription, the reactions were treated twice with DNAse I (New England Biolabs) and quenched with 10% volume of 0.5 M EDTA pH 8.0. The reactions were cleaned up with RNA Clean and Concentrator 100 columns (Zymo, referred to as “kit-purified” in text). Purity was assessed using a 6% Tris-Borate-Urea gel (Invitrogen).

For native purification of Arm B, the protocol for *in vitro* transcription of full-length SChLAP1 was adopted with the following modifications: 1) 10% DMSO was used rather than 5% DMSO, 2) Arm B was purified on an ENrich™ SEC 650 24 mL column (Bio-Rad), and 3) a smaller IVT scale was employed since a lower amount of RNA was needed for native gel electrophoresis. As above, the RNA was maintained at room temperature during this process. This sample is also referred to as “FPLC-purified” in text.

### Native gel electrophoresis

For divalent metal dependent experiments, Arm A or Arm B RNA were annealed in filtration buffer at approximately 50 nM. Annealing reactions were carried out at 20 µL aliquots. For snap-cooling, RNA was incubated at 95°C for 5 minutes and then submerged in ice for 15 minutes. For slow-cooling, RNA was incubated at 95°C for 5 minutes, cooled stepwise by 1°C/minute until 25°C was reached, then incubated at 25°C for 10 minutes followed by 4°C for 10 minutes. Divalent metals were spiked in from 20X concentrated stocks such that the desired final concentration of divalent metal was obtained. The samples were then incubated at 37°C for 30 minutes. Thereafter, 6 µL of the RNA was mixed with 6 µL 2X sample buffer (filtration buffer spiked with desired final divalent metal concentration, 10% glycerol, and 0.1% NP-40). 10 µL of this mix was added into wells of a 12-well 6% DNA retardation gel (Invitrogen) that was pre-running for at least 30 minutes in 0.5X TBE at 150V. The gels ran for 1 hour and 45 minutes (Arm B) or 1 hour and 15 minutes (Arm A, or gels where Arm A and Arm B ran together) before staining with Diamond Nucleic Acid Stain (Promega) for 20 minutes. Gels were imaged on an iBright 1500 imaging system (ThermoFisher scientific).

For KCl-dependent experiments, RNA was resuspended in HEPES-EDTA buffer (50 mM K-HEPES, pH 7.5, 100 µM EDTA), and KCl was spiked in from 20X concentrated stocks such that the final desired concentration was obtained. RNA was annealed (snap- or slow-cooled) as above and incubated at 37°C for 30 minutes for consistency with divalent metal experiments. The RNA was mixed with the same 2X sample buffer as above, with the exception that the KCl concentration was adjusted for each sample.

## DATA DEPOSITION

Unprocessed sequencing files and initial outputs of SHAPEMapper are available at Gene Expression Omnibus (GSE243328). SHAPE reactivities after normalization and 5NIA rescaling are provided in Supplementary Appendix 1.

## SUPPLEMENTARY MATERIAL

The supplementary material contains supplementary figures, tables, and text as referenced in the manuscript.

## ACKNOWLEDGEMENT

We would like to acknowledge all members of the Hargrove lab, past and present, for invaluable feedback and support. We also thank Bill Day, Michael Peterson, and İrem Altan for assistance with computation. In addition, we would also like to acknowledge Marek Zorawski for assistance with computation and cell culture. We also thank Whitehead Institute for Biomedical Research Genome Technology Core for DMS sample preparation and sequencing, as well as Silvi Rouskin and Fengrui Zhang for help with DMS data processing. We thank the Duke University School of Medicine for the use of the Sequencing and Genomic Technologies Shared Resource, which provided MiSeq service, as well as the Duke Compute Cluster for use of their cluster.

