## Supplementary Material for "Structural analysis of the lncRNA SChLAP1 reveals protein binding interfaces and a conformationally heterogenous retroviral insertion"

**Supplementary Text 1**

Pseudoknot prediction in SChLAP1

We additionally processed our SHAPE data for pseudoknot formation using SHAPEKnots (Hajdin et al. 2013). One long-range pseudoknot (nucleotides 429-434 and 599-604, 165 nucleotides between stems) was predicted from our *in vitro* data (Fig. S5A). A pseudoknot of similar 5′ coordinates was predicted in our *ex cellulo* data (nucleotides 416-420 and 644-648, 224 nucleotides between predicted stems) (Fig. S5A). However, for both predicted pseudoknots, the SHAPE reactivities of these predicted base-paired nucleotides was significantly increased in our *in cellulo* data and was thus incompatible with pseudoknot formation in cells (Fig. S5B and S5C). We thus did not constrain our any of our structure models herein to contain pseudoknots.

**A**

| ± | score | % div. | % del. | % ins. | query sequence | position in query- begin | end | (left) | C + | matching repeat | repeat class/family | -position in repeat- (left) begin | end end | begin (left) | linkage id/graphic |
|---|---|---|---|---|---|---|---|---|---|---|---|---|---|---|---|
| ± | 2699 | 4.3 | 0.0 | 0.0 | SChLAP1 | 1 | 322 | (1114) | + | LTR12C | LTR/ERV1 | 1258 | 1579 | (0) | 1 |
| ± | 16 | 0.0 | 0.0 | 0.0 | SChLAP1 | 1088 | 1104 | (332) | + | (A)n | Simple_repeat | 1 | 17 | (0) | 2 |
| ± | 1973 | 11.2 | 3.2 | 3.5 | SChLAP1 | 1123 | 1436 | (0) | + | THE1B | LTR/ERVL-MaLR | 1 | 313 | (51) | 3 |

**B**

```
SChLAP1                 1 GCTTTTATGAGCTGTAACACTCACCGCGAAGGTCCGCAGCTTCACTCCTG 50
                            i                               i
LTR12C#LTR/ER        1258 GCCTTTATGAGCTGTAACACTCACCGCGAAGGTCTGCAGCTTCACTCCTG 1307

SChLAP1                51 AAGCCAGCGAGACCACGAGCCTACTGGGAGGAACGAACAACTCCCGACGC 100
                                               i  i                   v
LTR12C#LTR/ER        1308 AAGCCAGCGAGACCACGAGCCCACCGGGAGGAACGAACAACTCCAGACGC 1357

SChLAP1               101 GCCGCCTTAAGAGCTGTAACACTCACCGCGAAGGTCTGCAGCTTCACTCC 150

LTR12C#LTR/ER        1358 GCCGCCTTAAGAGCTGTAACACTCACCGCGAAGGTCTGCAGCTTCACTCC 1407

SChLAP1               151 TGAGCCAGCGAGACCACGAACCCACCAGAAGGAAAAACTCCGAACACAT 200
                                                            i
LTR12C#LTR/ER        1408 TGAGCCAGCGAGACCACGAACCCACCAGAAGGAAGAAACTCCGAACACAT 1457

SChLAP1               201 CTGAACATCAGAAGCAACAAACTCCGGACACGCCGCCTTTAAGAACTGTA 250
                           i            v              i              i
LTR12C#LTR/ER        1458 CCGAACATCAGAAGGAACAAACTCCGGACGCGCCGCCTTTAAGAGCTGTA 1507

SChLAP1               251 ACACTCACTGCGAGGGTCCGCGGCTTCATTCTTGAAGTGAGTGAGACCAA 300
                                  i                             v
LTR12C#LTR/ER        1508 ACACTCACCGCGAGGGTCCGCGGCTTCATTCTTGAAGTCAGTGAGACCAA 1557

SChLAP1               301 GAACCCACCAGTTCTGGACACA 322
                                    i   i
LTR12C#LTR/ER        1558 GAACCCACCAATTCCGGACACA 1579
```

**C**

```
SChLAP1              1123 TGATATGGTTTGGCTGTGTCCCCACCCAAATATCATCTTGAATTGTAGCT 1172
                                                         v
THE1B#LTR/ERV           1 TGATATGGTTTGGCTGTGTCCCCACCCAAATCTCATCTTGAATTGTAGCT 50

SChLAP1              1173 CCCATAATTCCCACGTGTTGTGGGAGGGACCCGGTGGGAGATAATTGTAT 1222
                                            i                     i      v
THE1B#LTR/ERV          51 CCCATAATTCCCACGTGTCGTGGGAGGGACCCGGTGGGAGGTAATTGAAT 100

SChLAP1              1223 CATGGGGGTGGTTC--CCCCATACTATTCTCATAGTAGTGAATAAGTCTC 1270
                                  i  v  --i   i i  i     i ii
THE1B#LTR/ERV         101 CATGGGGGCGGGTCTTTCCCGTGCTGTTCTCGTGATAGTGAATAAGTCTC 150

SChLAP1              1271 ACAAAATCTGATGGTTTTATGAGGGAAAACCCCTTTCACCTGGTTCTCAT 1320
                            i i               i i  ii i-----   v    --- v  ?
THE1B#LTR/ERV         151 ACGAGATCTGATGGTTTTATAAAGGGGAG-----TTCCCCTG---CACAN 192

SChLAP1              1321 TCTCTTCTCTGGTCTGTCGTCATGTAAGACATGCCTTT----CACCTTC- 1365
                          v ---     v i   i  i          i  ?    ---- v     -
THE1B#LTR/ERV         193 GC---TCTCTTGCCTGCCGCCATGTAAGACGTGNCTTTGCTCCTCCTTCG 239

SChLAP1              1366 ---TCCACCATGACTGTGAGGCCTCCCCAGCCACGTGGAACTGTGAGCCC 1412
                          ---   i      i                                 i
THE1B#LTR/ERV         240 CCTTCCGCCATGATTGTGAGGCCTCCCCAGCCACGTGGAACTGTGAGTCC 289

SChLAP1              1413 ATTAAACCTCTTTCACTTATAAAT 1436
                                        vi
THE1B#LTR/ERV         290 ATTAAACCTCTTTCCTTTATAAAT 313
```

Figure S1. RepeatMasker analysis of SChLAP1 Iso. 1. A) Complete results table of RepeatMasker analysis of SChLAP1 (condensed version shown in Fig. 1B). B) LTR12C alignment of SChLAP1. C) THE1B alignment of SChLAP1.

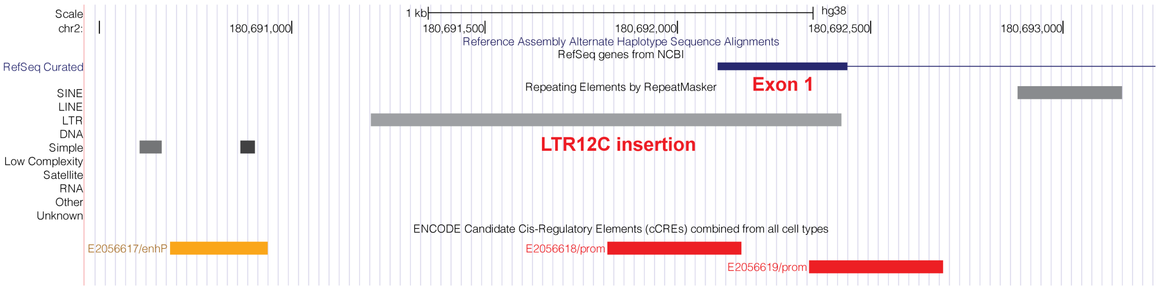

Figure S2. UCSC genome browser (Kent et al. 2002) display of SChLAP1 promoter and exon 1. Position of LTR12C insertion and exon 1 is indicated.

A

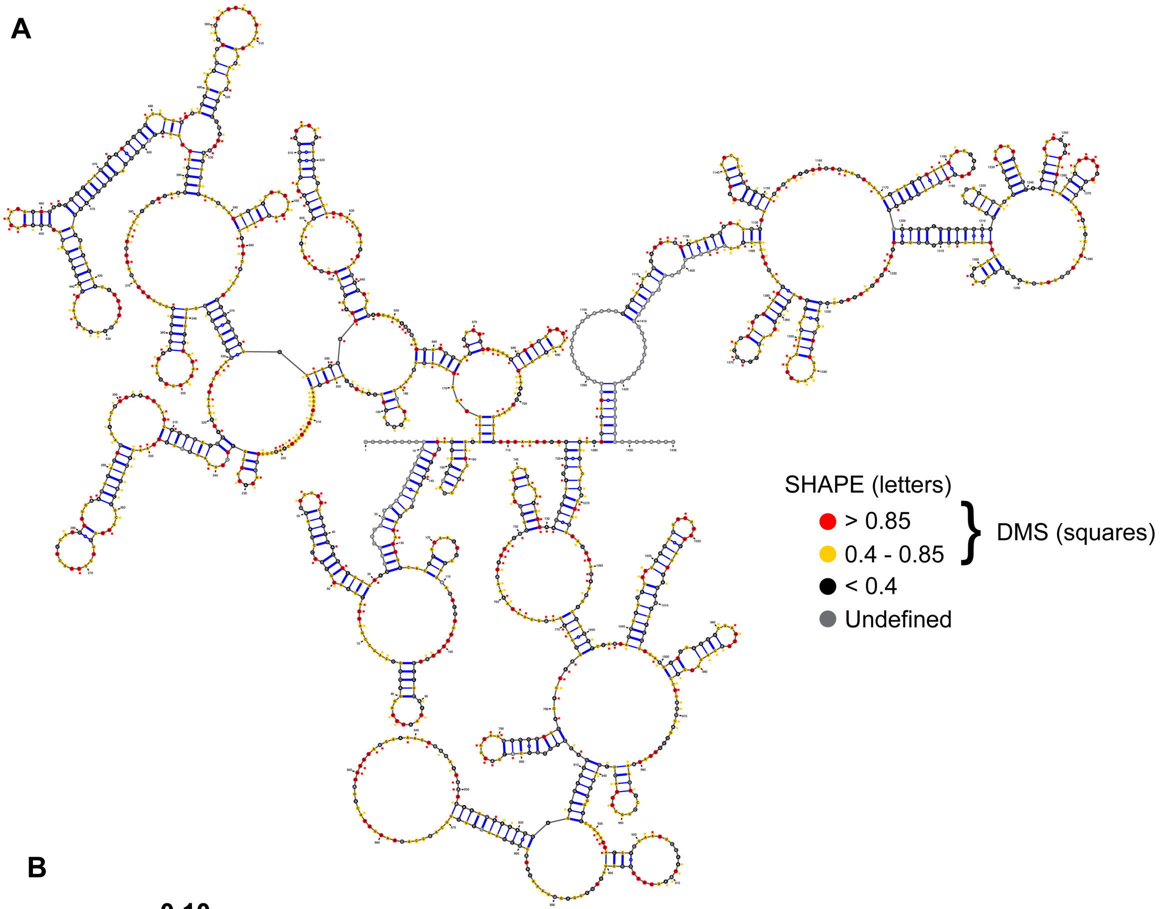

B

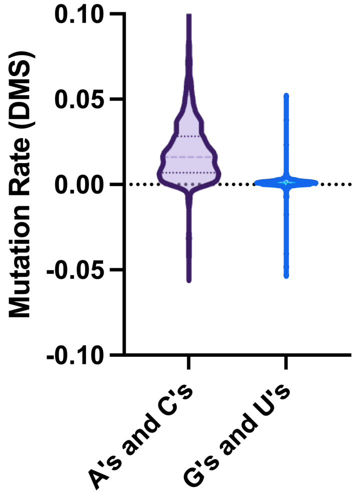

Figure S3. MFE structure model of *in vitro* SChLAP1. A) SHAPE-informed MFE structure of *in vitro* SChLAP1 generated through semi-native purification. DMS reactivities are overlaid over SHAPE-informed structure. MFE structure was generated in RNAstructure (Reuter and Mathews 2010) and visualized in VARNA (Darty et al. 2009). Arc diagram for *in vitro* data shown in Fig. S4.

B) Mutation rate of adenosines (A's) and cytidines (C's) versus guanosines (G's) and uridines (U's) from DMS probing.

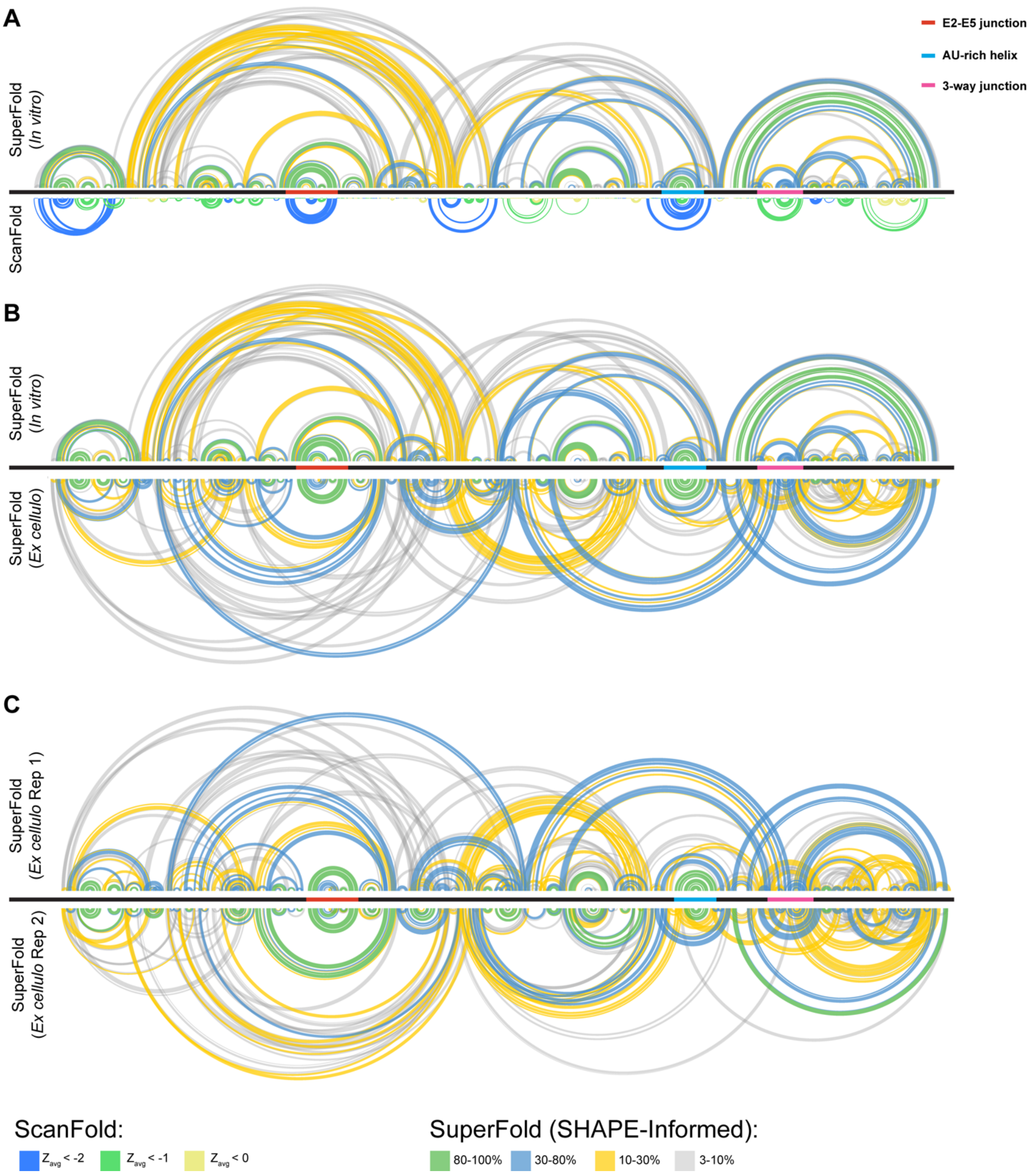

Figure S4. Comparison of arc diagrams between A) *in vitro* SHAPE-probed SChLAP1 and ScanFold-prediction, B) *in vitro* SHAPE probed and representative replicate of *ex cellulo* SHAPE probing (same as Fig. 3), and C) both replicates of *ex cellulo* SHAPE probing. Positions of E2-E5 junction, AU-rich helix, and 3WJ are indicated with colored bars. Figure generated in Integrative Genomics Viewer (Robinson et al. 2011).

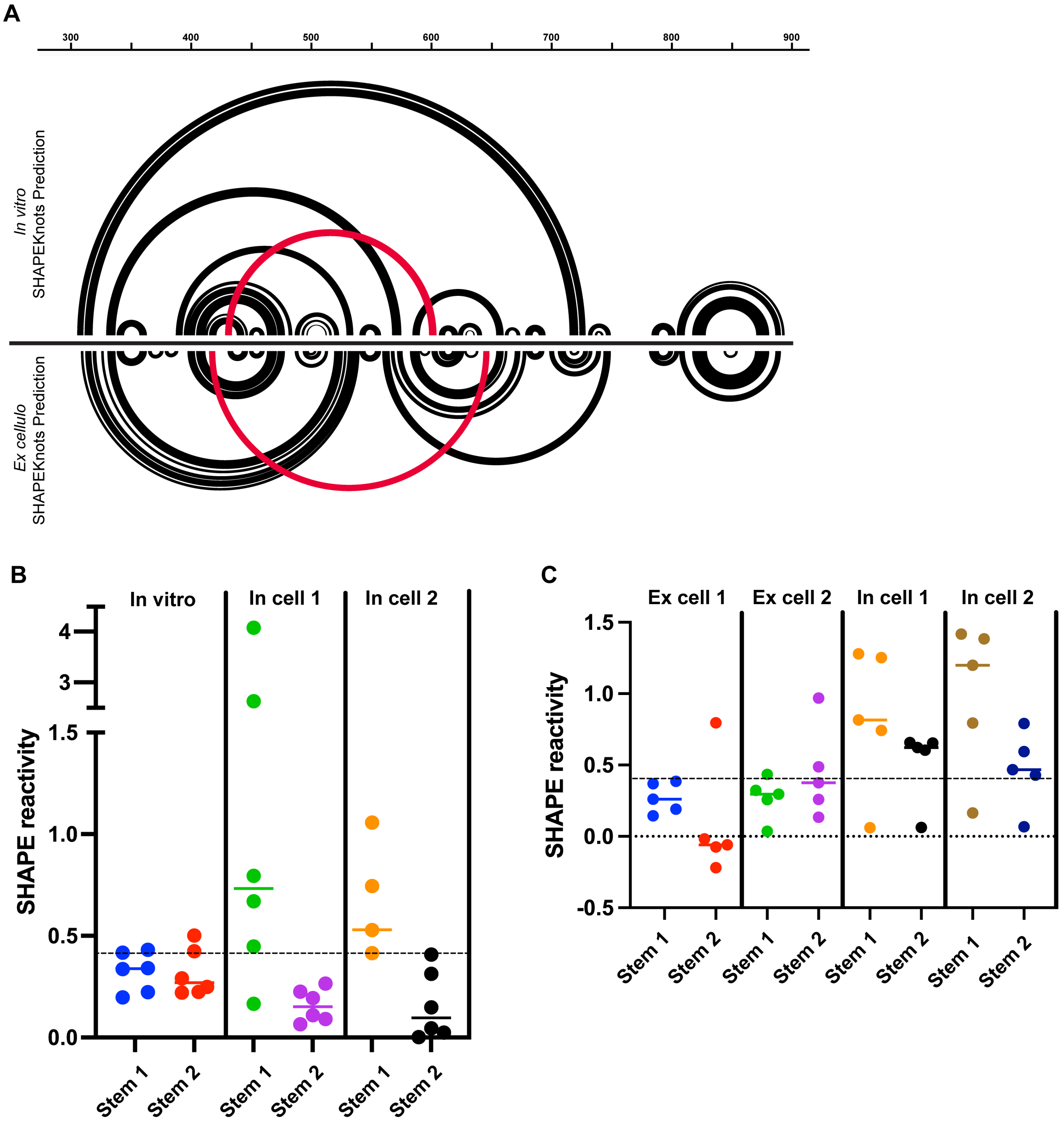

Figure S5. SHAPEknots analysis of SChLAP1. A) Representative arc diagrams of SChLAP1 prediction window generated through SHAPEknots (Hajdin et al. 2013). Red arcs denote predicted pseudoknots in *in vitro* and *ex cellulo* datasets. Figure generated in IGV (Robinson et al. 2011). B) SHAPE reactivities for the *in vitro*-predicted pseudoknot and the same nucleotides in *in cellulo* replicates 1 and 2, divided by stem of the predicted pseudoknot. Colored lines denote median. A line at SHAPE = 0.4 is drawn for reference as this reactivity is where base-pairing is no longer favorable (Hajdin et al. 2013). For *in cellulo* replicate 2, stem 1, one nucleotide of undefined reactivity was not plotted, and a second nucleotide with outlier SHAPE reactivity (see Methods) was also not plotted, although it was maintained for SHAPEknots prediction and calculation of the median. C) SHAPE reactivities for the *ex cellulo* predicted pseudoknot for both *ex cellulo* and *in cellulo* replicates, divided by putative stem as in panel B.

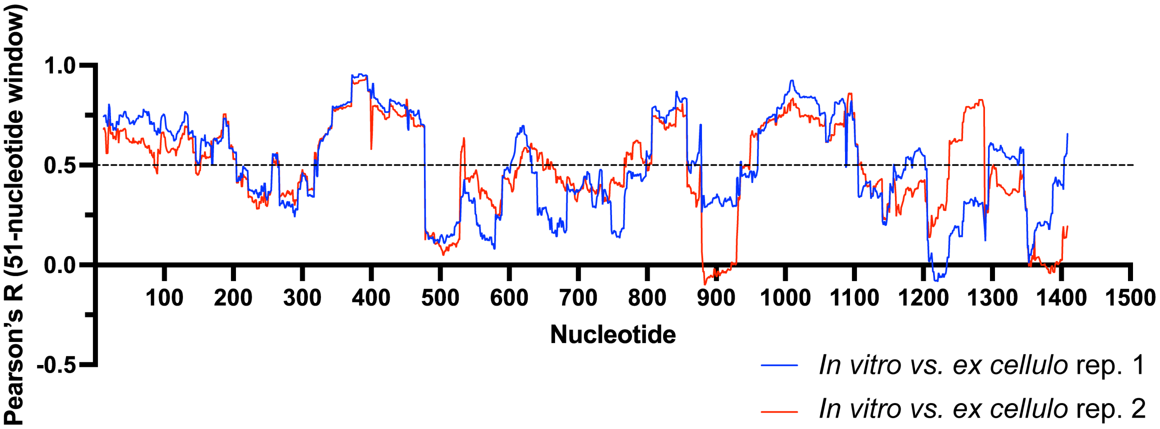

Figure S6. Correlation between *ex cellulo* and *in vitro* SChLAP1. Pearson correlation coefficients were calculated in 51-nucleotide sliding windows (25 nucleotides on each side of a central nucleotide). Correlation windows were truncated at the 5′ and 3′ ends until no fewer than 10 nucleotides were included in the calculation. A dashed line at R = 0.5 is drawn for reference.

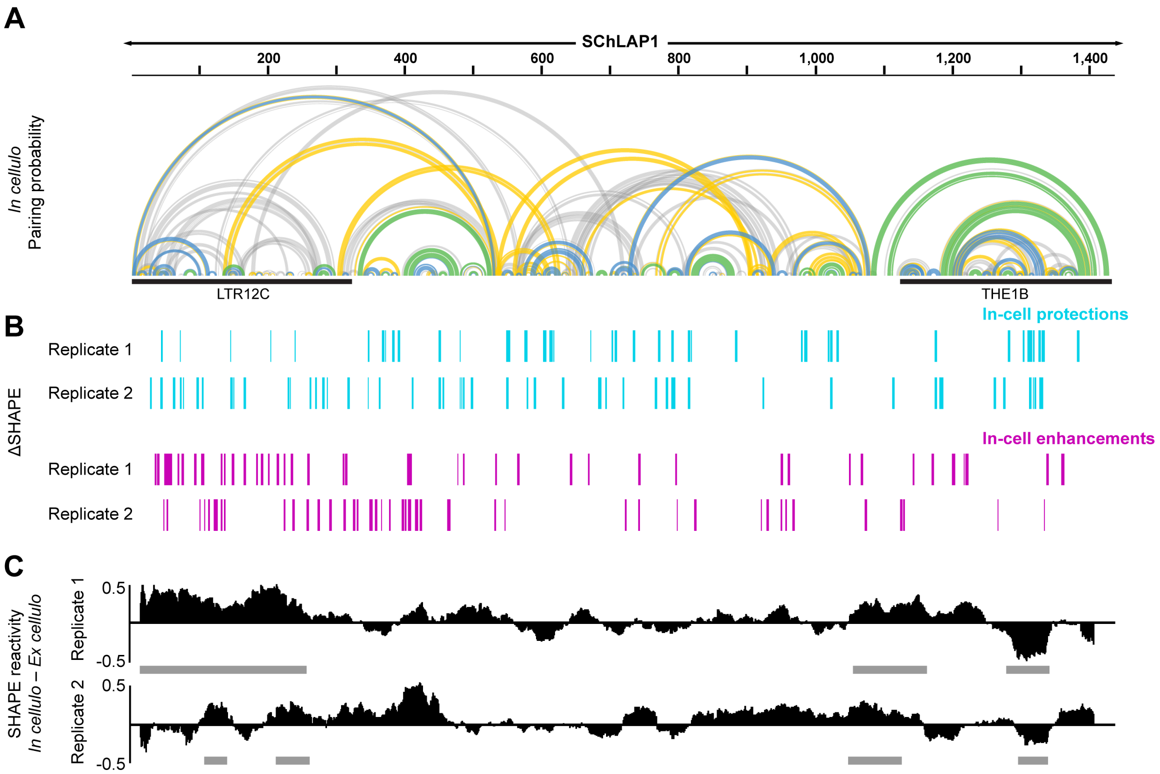

Figure S7. *In cellulo* SHAPE and comparison to *ex cellulo* models. A) Arc diagram of *in cellulo* probed SChLAP1, representative of two biological replicates. Color scheme is the same as Fig. 3A) Location of in-cell protections and in-cell enhancements from each replicate of ΔSHAPE analysis. Overlapping regions between replicates were merged and displayed in Figure 4 (see Methods). C) Difference plot of SHAPE reactivities between each replicate of *in cellulo* and *ex cellulo* reactivities. Reactivities were smoothened over 51-nucleotide windows (see Methods). Positive values indicate regions that are more reactive *in cellulo*. Regions of significant change (see Methods) overlapping between both replicates are indicated by grey bars. Figure generated in Integrative Genomics Viewer (Robinson et al. 2011).

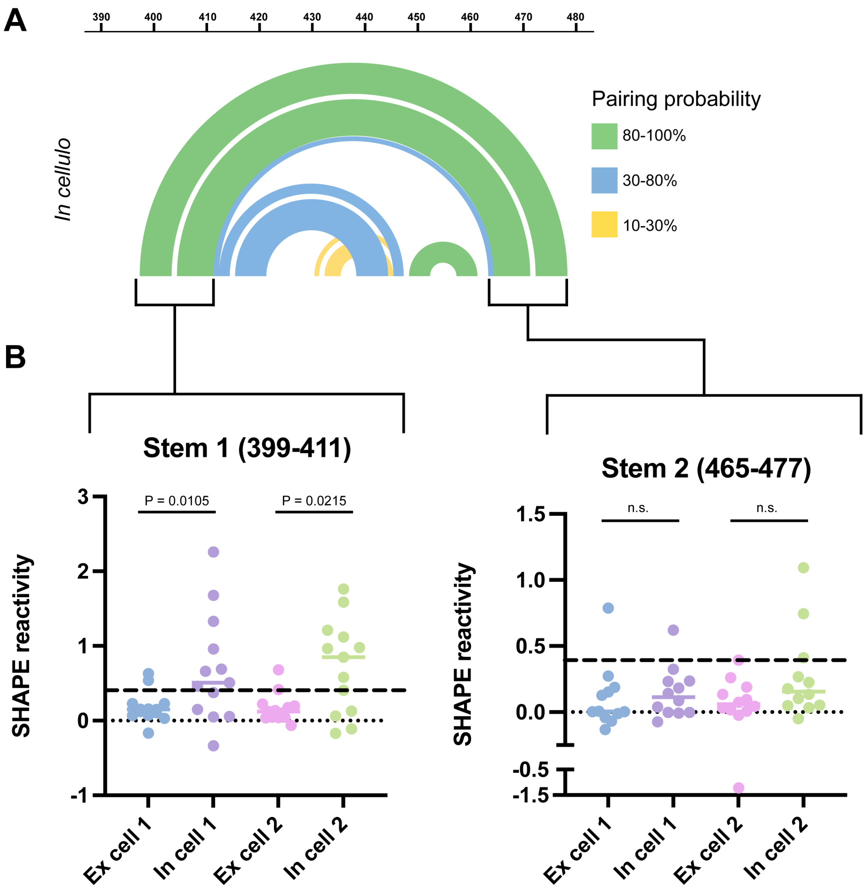

Figure S8. Analysis of SHAPE reactivities of the E2-E5 junction in cells. A) SHAPE-informed arc diagram of the E2-E5 junction (nucleotides 398-478) from a representative replicate of *in cellulo* probing. Pairing probabilities less than 10% are not depicted. B) Analysis of SHAPE reactivities of the largest stem within in this structure. Nucleotides 398, 412, 464, and 478 are ignored in this analysis as they are located at the end of helices and thus may have higher SHAPE reactivities, i.e. only internal positions were considered. Statistical comparisons were performed with a Wilcoxon t-test (comparing *ex cellulo* 1 with *in cellulo* 1 and *ex cellulo* 2 with *in cellulo* 2, as was performed with ΔSHAPE) with a cutoff of 0.05 for significance.

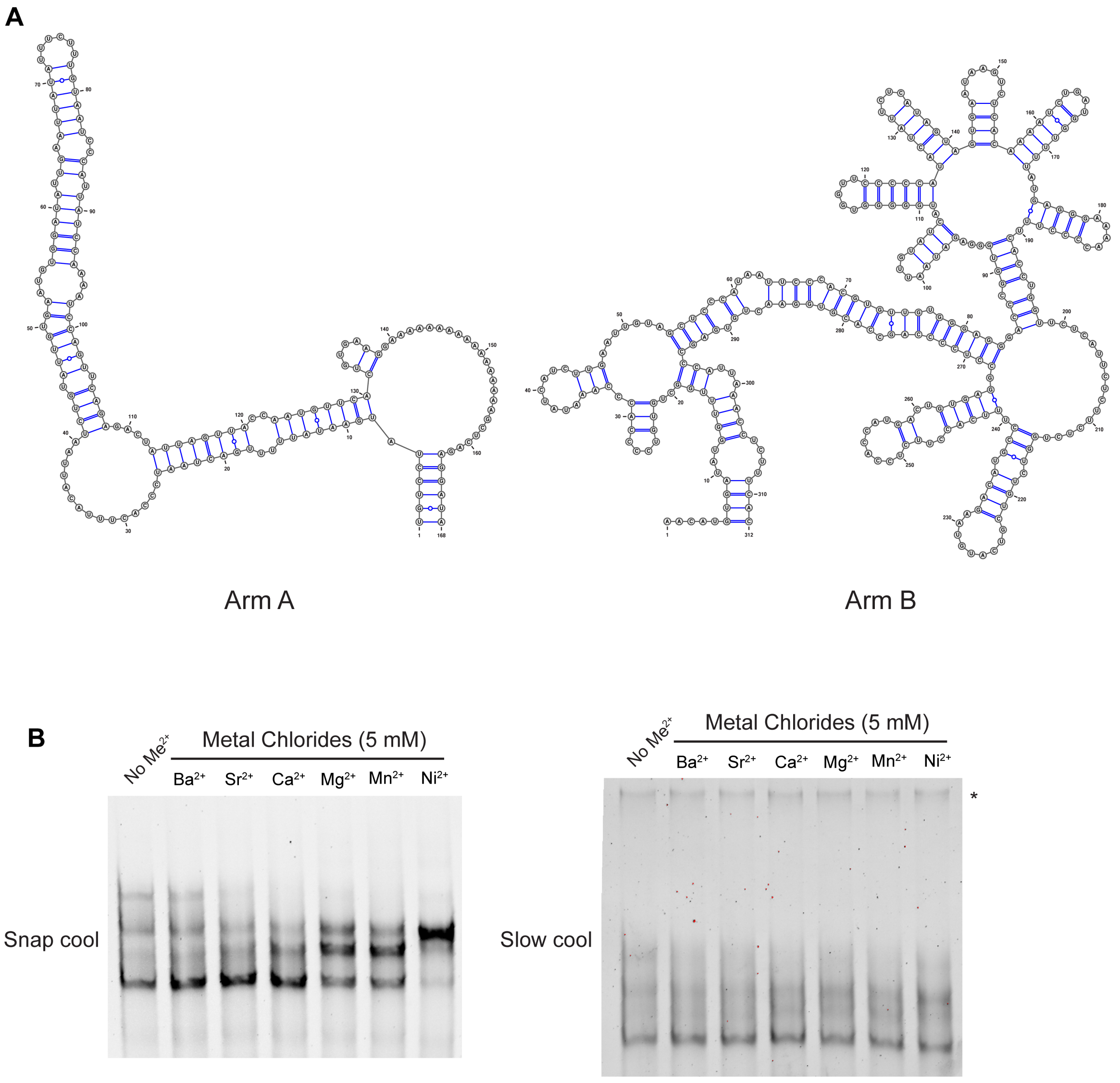

Figure S9. Characterization of Arm A and Arm B by native gel electrophoresis. A) MFE structures of Arm A (left) and Arm B (right) calculated in RNAstructure (Bellaousov et al. 2013) and visualized in VARNA (Darty et al. 2009). B) Native gel electrophoresis of kit-purified Arm B that was either snap cooled (left) or slow cooled (right) and thereafter incubated with the given divalent cations. Arm B was generated through the "kit purified" method. Each gel was performed in triplicate. Asterisks denote dimer, which was typically favored in slow cooling.

**A**

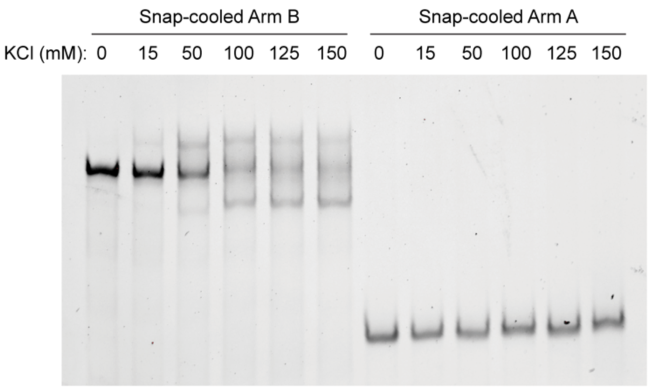

**B**

Maximum length = 30 nucleotides

| Position | Length | QGRS |
|---|---|---|
| 449 | 30 | GGACACAAGGUCCACCUUCCAGGAAUAUGG |
| 638 | 19 | GGAGCGGGAUGGAGAAAGG |
| 719 | 30 | GGAGAUGGUGAACCCAAGAGGCUCUGCUGG |
| 1195 | 16 | GGAGGGACCCGGUGGG |

Maximum length = 45 nucleotides

| Position | Length | QGRS |
|---|---|---|
| 447 | 32 | GGGGACACAAGGUCCACCUUCCAGGAAUAUGG |
| 638 | 19 | GGAGCGGGAUGGAGAAAGG |
| 719 | 30 | GGAGAUGGUGAACCCAAGAGGCUCUGCUGG |
| 786 | 40 | GGUGAAUAUCAGCACUGGGACCAGUUAGCAGAGGAAAAGG |
| 1194 | 35 | GGGAGGGACCCGGUGGGAGAUAAUUGUAUCAUGGG |

**C**

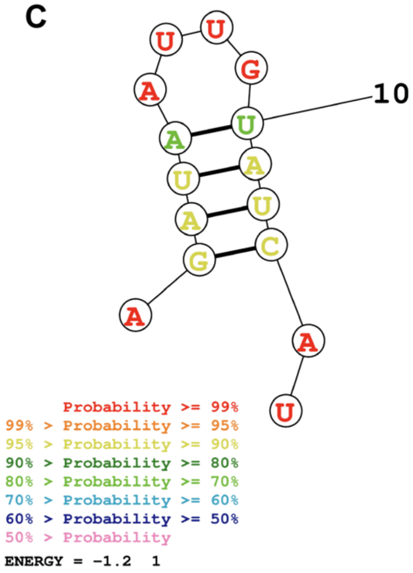

Figure S10. G-quadruplex analysis of Arm B. A) Native gel electrophoresis of kit-purified Arm A and Arm B in varying KCl concentrations. Gel is representative of three biological replicates. B) QGRS Mapper (Kikin et al. 2006) prediction of SChLAP1 with 30 nucleotides (left) and 45

nucleotides (right) set as the maximum allowed G-quadruplex length. Blue letters denote putative G-quadruplex forming regions, and green letters denote predicted hairpin shown in Panel C. C) Structure of predicted hairpin generated in RNAstructure (Bellaousov et al. 2013).

Table S1. SChLAP1 primate accession codes

| Name | Organism | Common Name | Accession Number |
|---|---|---|---|
| SChLAP1 isoform 1 | *Homo sapiens* | Human | NR_104319.1 |
| SChLAP1 isoform 2 | *Homo sapiens* | Human | NR_104320.1 |
| SChLAP1 isoform 3 | *Homo sapiens* | Human | NR_104321.1 |
| SChLAP1 isoform 4 | *Homo sapiens* | Human | NR_104322.1 |
| SChLAP1 isoform 5 | *Homo sapiens* | Human | NR_104323.1 |
| SChLAP1 isoform 6 | *Homo sapiens* | Human | NR_104324.1 |
| SChLAP1 isoform 7 | *Homo sapiens* | Human | NR_104325.1 |
| PREDICTED: Cercocebus atys uncharacterized LOC105579891 (LOC105579891), ncRNA | *Cercocebus atys* | Sooty Mangabey | XR_001012906.1 |
| PREDICTED: Mandrillus leucophaeus uncharacterized LOC105529207 (LOC105529207), ncRNA | *Mandrillus leucophaeus* | Drill | XR_001004474.1 |
| PREDICTED: Colobus angolensis palliatus uncharacterized LOC105524242 (LOC105524242), ncRNA | *Colobus angolensis palliatus* | Angolan Colobus | XR_001003724.1 |
| PREDICTED: Pan troglodytes uncharacterized LOC107973853 (LOC107973853), ncRNA | *Pan troglodytes* | Chimpanzee | XR_001716344.3 |
| PREDICTED: Hylobates moloch uncharacterized LOC116809510 (LOC116809510), ncRNA | *Hylobates moloch* | Silvery Gibbon | XR_004369869.1 |
| PREDICTED: Nomascus leucogenys uncharacterized | *Nomascus leucogenys* | Northern White-Cheeked Gibbon | XR_001113961.2 |

| Name | Organism | Common Name | Accession Number |
| --- | --- | --- | --- |
| LOC105738214 (LOC105738214), ncRNA | | | |
| PREDICTED: Pan paniscus uncharacterized LOC130541288 (LOC130541288), ncRNA | *Pan paniscus* | Pygmy Chimpanzee | XR_008955326.1 |
| PREDICTED: Gorilla gorilla gorilla uncharacterized LOC109025949 (LOC109025949), ncRNA | *Gorilla gorilla gorilla* | Western Lowland Gorilla | XR_008677752.1 |

Table S2. All DNA sequences.

| Sequence Name | Sequence (5'→3'; Sense Strand) |
| --- | --- |
| SChLAP1 Iso. 1 template | GCTTTTATGAGCTGTAACACTCACCGCGAAGGTCCGCAGCTTCA<br>CTCCTGAAGCCAGCGAGACCACGAGCCTACTGGGAGGAACGAA<br>CAACTCCCGACGCGCCGCCTTAAGAGCTGTAACACTCACCGCG<br>AAGGTCTGCAGCTTCACTCCTGAGCCAGCGAGACCACGAACCC<br>ACCAGAAGGAAAAAACTCCGAACACATCTGAACATCAGAAGCAA<br>CAAACTCCGGACACGCCGCCTTTAAGAACTGTAACACTCACTGC<br>GAGGGTCCGCGGCTTCATTCTTGAAGTGAGTGAGACCAAGAAC<br>CCACCAGTTCTGGACACAATTTCAAGTCCTCAGGTGCCATCAAT<br>ATTCTGAAAATGGCAGTGATTTTTATTCAACCTGTATAAGGCACT<br>TTCACCATGTACCTGGAAGCAACATCTACATCTTTTTCAGCAATC<br>TAGATGCTGGGGACACAAGGTCCACCTTCCAGGAATATGGCCA<br>TGACACCAGAAATCACAAACATGATGAGAATGGAATGACTGGGG<br>AAGAAGTGCCAGATGCTTCACTTGTAAATGAAGACCCAGCCTCT<br>GGGGATGCAGATACCACCTCCCTGAAGAAGCTGAATATCTGCA<br>GATAAGTGGAGTTCACCAATGATGAGGAGCGGGATGGAGAAAG<br>GAGGTAGGGAGAGTCATCCAAGGAACATGAGCAACATGTTAAA<br>AGCCAAGTGGTTTAATTTCTGGAGATGGTGAACCCAAGAGGCTC<br>TGCTGGGAGACAACAAAAATAATGAAGAATTGAACCAGAGTCCG<br>GTGAATATCAGCACTGGGACCAGTTAGCAGAGGAAAAGGAAAG<br>AATAAAAGCGAAAAGAATGAAGAGTCATATGATTACCAACTTTTC<br>CTTTTTCATATAAATTGAGTGTATATGGGTCTGGAACAACCTGAA<br>TTTCCATCAAGTCCTGGCTAACCTCATTATGTCCTATGAATATTT<br>TTGACTAATCCCACTTTACATTAATCTGTATTGTGAATGTGGATA<br>TTGAATTATATTTCTTTGTAATCCCATTATCCAAAATCCAGTTCAG<br>AGACTATTAGTTACCAATGTTCACTGTGAAGGAAAAAAAAAAAAA<br>AAAAGCTCAGAGGATAAACATGTGATATGGTTTGGCTGTGTCCC<br>CACCCAAATATCATCTTGAATTGTAGCTCCCATAATTCCCACGTG<br>TTGTGGGAGGGACCCGGTGGGAGATAATTGTATCATGGGGGTG |

| Sequence Name | Sequence (5'→3'; Sense Strand) |
|---|---|
| | GTTCCCCCATACTATTCTCATAGTAGTGAATAAGTCTCACAAAAT CTGATGGTTTTATGAGGGAAAACCCCTTTCACCTGGTTCTCATT CTCTTCTCTGGTCTGTCGTCATGTAAGACATGCCTTTCACCTTCT CCACCATGACTGTGAGGCCTCCCCAGCCACGTGGAACTGTGAG CCCATTAAACCTCTTTCACTTATAAAT |
| SChLAP1 Arm A (949-1116) | TGTCCTATGAATATTTTTGACTAATCCCACTTTACATTAATCTGTA TTGTGAATGTGGATATTGAATTATATTTCTTTGTAATCCCATTATC CAAAATCCAGTTCAGAGACTATTAGTTACCAATGTTCACTGTGAA GGAAAAAAAAAAAAAAAAAGCTCAGAGGATA |
| SChLAP1 Arm B (1117-1428) | AACATGTGATATGGTTTGGCTGTGTCCCCACCCAAATATCATCTT GAATTGTAGCTCCCATAATTCCCACGTGTTGTGGGAGGGACCC GGTGGGAGATAATTGTATCATGGGGGTGGTTCCCCCATACTATT CTCATAGTAGTGAATAAGTCTCACAAAATCTGATGGTTTTATGAG GGAAAACCCCTTTCACCTGGTTCTCATTCTCTTCTCTGGTCTGTC GTCATGTAAGACATGCCTTTCACCTTCTCCACCATGACTGTGAG GCCTCCCCAGCCACGTGGAACTGTGAGCCCATTAAACCTCTTTC AC |
| SChLAP1 Iso. 1 Amplicon 1 | GCTTTTATGAGCTGTAACACTCACCGCGAAGGTCCGCAGCTTCA CTCCTGAAGCCAGCGAGACCACGAGCCTACTGGGAGGAACGAA CAACTCCCGACGCGCCGCCTTAAGAGCTGTAACACTCACCGCG AAGGTCTGCAGCTTCACTCCTGAGCCAGCGAGACCACGAACCC ACCAGAAGGAAAAAACTCCGAACACATCTGAACATCAGAAGCAA CAAACTCCGGACACGCCGCCTTTAAGAACTGTAACACTCACTGC GAGGGTCCGCGGCTTCATTCTTGAAGTGAGTGAGACCAAGAAC CCACCAGTTCTGGACACAATTTCAAGTCCTCAGGTGCCATCAAT ATTCTGAAAATGGCAGTGATTTTTATTCAACCTGTATAAGGCACT TTCACCATGTACCTGGAAGCAACATCTACATCTTTTTCAGCAATC TAGATGCTGGGGACACAAGGTCCACCTTCCAGGAATATGGCCA TGACACCAGAAATCACAAA |
| SChLAP1 Iso. 1 Amplicon 2 | TGTACCTGGAAGCAACATCTACATCTTTTTCAGCAATCTAGATGC TGGGGACACAAGGTCCACCTTCCAGGAATATGGCCATGACACC AGAAATCACAAACATGATGAGAATGGAATGACTGGGGAAGAAGT GCCAGATGCTTCACTTGTAAATGAAGACCCAGCCTCTGGGGAT GCAGATACCACCTCCCTGAAGAAGCTGAATATCTGCAGATAAGT GGAGTTCACCAATGATGAGGAGCGGGATGGAGAAAGGAGGTAG GGAGAGTCATCCAAGGAACATGAGCAACATGTTAAAAGCCAAGT GGTTTAATTTCTGGAGATGGTGAACCCAAGAGGCTCTGCTGGG AGACAACAAAAATAATGAAGAATTGAACCAGAGTCCGGTGAATA TCAGCACTGGGACCAGTTAGCAGAGGAAAAGGAAAGAATAAAA GCGAAAAGAATGAAGAGTCATATGATTACCAACTTTTCCTTTTTC ATATAAATTGAGTGTATATGGG |

| Sequence Name | Sequence (5'→3'; Sense Strand) |
| --- | --- |
| SChLAP1 Iso. 1 Amplicon 3 | CTGGGACCAGTTAGCAGAGGAAAAGGAAAGAATAAAAGCGAAAAGAATGAAGAGTCATATGATTACCAACTTTTCCTTTTTCATATAAATTGAGTGTATATGGGTCTGGAACAACCTGAATTTCCATCAAGTCCTGGCTAACCTCATTATGTCCTATGAATATTTTTGACTAATCCCACTTTACATTAATCTGTATTGTGAATGTGGATATTGAATTATATTTCTTTGTAATCCCATTATCCAAAATCCAGTTCAGAGACTATTAGTTACCAATGTTCACTGTGAAGGAAAAAAAAAAAAAAAAAAGCTCAGAGGATAAACATGTGATATGGTTTGGCTGTGTCCCCACCCAAATATCATCTTGAATTGTAGCTCCCATAATTCCCACGTGTTGTGGGAGGGACCCGGTGGGAGATAATTGTATCATGGGGGTGGTTCCCCCATACTATTCTCATAGTAGTGAATAAGTCTCACAAAATCTGATGGTTTTATGAGGGAAAAC |
| SChLAP1 Iso. 1 Amplicon 4 | GACCCGGTGGGAGATAATTGTATCATGGGGGTGGTTCCCCCATACTATTCTCATAGTAGTGAATAAGTCTCACAAAATCTGATGGTTTTATGAGGGAAAACCCCTTTCACCTGGTTCTCATTCTCTTCTCTGGTCTGTCGTCATGTAAGACATGCCTTTCACCTTCTCCACCATGACTGTGAGGCCTCCCCAGCCACGTGGAACTGTGAGCCCATTAAACCTCTTTCACTTATAAAT |

Table S3. All primer sequences for PCR and/or RT. [mX] indicates nucleotide with a 2'–OMe modification. N indicates randomization of A, C, G, or T.

| Sequence Name | Forward Primer Sequence (5'→3') | Reverse Primer Sequence (5'→3') |
|---|---|---|
| SChLAP1 Iso. 1 | CTAATACGACTCACTATAGCTTTTATGAGCTGTAACACTCACCGC | [mA][mU]TTATAAGTGAAAGAGGTTTAATGGGCTCACAGTTCC |
| Arm A | GAAATTAATACGACTCACTATAGTGTCCTATGAATATTTTTGACTAATCCCAC | [mU][mA]TCCTCTGAGCTTTTTTTTTTTTTTTTTCCTTCAC |
| Arm B | GAAATTAATACGACTCACTATAGAACATGTGATATGGTTTGGCTGTGTC | [mG][mU]GAAAGAGGTTTAATGGGCTCACAGTTC |
| SChLAP1 Iso. 1 Amplicon 1 Blunt end | GCTTTTATGAGCTGTAACACTCACCGC | TTTGTGATTTCTGGTGTCATGGCCATATTCC |
| SChLAP1 Iso. 1 Amplicon 2 Blunt end | ATGTACCTGGAAGCAACATCTACATCTTTTTCAGC | CCCATATACACTCAATTTATATGAAAAAGGAAAAGTTGGTAATCATATGACTC |
| SChLAP1 Iso. 1 Amplicon 3 Blunt end | CTGGGACCAGTTAGCAGAGGAAAAGG | GTTTTCCCTCATAAAACCATCAGATTTTGTGAGACTTA |
| SChLAP1 Iso. 1 Amplicon 4 Blunt end | GACCCGGTGGGAGATAATTGTATCATG | ATTTATAAGTGAAAGAGGTTTAATGGGCTCACAGTTCC |
| SChLAP1 Iso. 1 Amplicon 1 PCR1 | CCCTACACGACGCTCTTCCGATCTNNNNNGCTTTTATGAGCTGTAACACTCACCGC | GACTGGAGTTCAGACGTGTGCTCTTCCGATCTNNNNNTTTGTGATTTCTGGTGTCATGGCCATATTCC |
| SChLAP1 Iso. 1 Amplicon 2 PCR1 | CCCTACACGACGCTCTTCCGATCTNNNNNATGTACCTGGAAGCAACATCTACATCTTTTTCAGC | GACTGGAGTTCAGACGTGTGCTCTTCCGATCTNNNNNCCCATATACACTCAATTTATATGAAAAAGGAAAAGTTGGTAATCATATGACTC |

| Sequence Name | Forward Primer Sequence (5'→3') | Reverse Primer Sequence (5'→3') |
|---|---|---|
| SChLAP1 Iso. 1 Amplicon 3 PCR1 | CCCTACACGACGCTCTTCCGATCTNNNNNCTGGGACCAGTTAGCAGAGGAAAAGG | GACTGGAGTTCAGACGTGTGCTCTTCCGATCTNNNNNGTTTTCCCTCATAAAACCATCAGATTTTGTGAGACTTA |
| SChLAP1 Iso. 1 Amplicon 4 PCR1 | CCCTACACGACGCTCTTCCGATCTNNNNNGACCCGGTGGGAGATAATTGTATCATG | GACTGGAGTTCAGACGTGTGCTCTTCCGATCTNNNNNATTTATAAGTGAAAGAGGTTTAATGGGCTCACAGTTCC |

Table S4. RNA sequences made by *in vitro* transcription.

| Sequence Name | Sequence (5'→3') |
|---|---|
| SChLAP1 Iso. 1 | GCUUUUAUGAGCUGUAACACUCACCGCGAAGGUCCGCAG<br>CUUCACUCCUGAAGCCAGCGAGACCACGAGCCUACUGGG<br>AGGAACGAACAACUCCCGACGCGCCGCCUUAAGAGCUGU<br>AACACUCACCGCGAAGGUCUGCAGCUUCACUCCUGAGCC<br>AGCGAGACCACGAACCCACCAGAAGGAAAAAACUCCGAAC<br>ACAUCUGAACAUCAGAAGCAACAAACUCCGGACACGCCGC<br>CUUUAAGAACUGUAACACUCACUGCGAGGGUCCGCGGCU<br>UCAUUCUUGAAGUGAGUGAGACCAAGAACCCACCAGUUC<br>UGGACACAAUUUCAAGUCCUCAGGUGCCAUCAAUAUUCU<br>GAAAAUGGCAGUGAUUUUUAUUCAACCUGUAUAAGGCAC<br>UUUCACCAUGUACCUGGAAGCAACAUCUACAUCUUUUUC<br>AGCAAUCUAGAUGCUGGGGACACAAGGUCCACCUUCCAG<br>GAAUAUGGCCAUGACACCAGAAAUCACAAACAUGAUGAGA<br>AUGGAAUGACUGGGGAAGAAGUGCCAGAUGCUUCACUUG<br>UAAAUGAAGACCCAGCCUCUGGGGAUGCAGAUACCACCU<br>CCCUGAAGAAGCUGAAUAUCUGCAGAUAAGUGGAGUUCA<br>CCAAUGAUGAGGAGCGGGAUGGAGAAAGGAGGUAGGGA<br>GAGUCAUCCAAGGAACAUGAGCAACAUGUUAAAAGCCAAG<br>UGGUUUAAUUUCUGGAGAUGGUGAACCCAAGAGGCUCUG<br>CUGGGAGACAACAAAAAUAAUGAAGAAUUGAACCAGAGUC<br>CGGUGAAUAUCAGCACUGGGACCAGUUAGCAGAGGAAAA<br>GGAAAGAAUAAAAGCGAAAAGAAUGAAGAGUCAUAUGAUU<br>ACCAACUUUUCCUUUUUCAUAUAAAUUGAGUGUAUAUGG<br>GUCUGGAACAACCUGAAUUUCCAUCAAGUCCUGGCUAAC<br>CUCAUUAUGUCCUAUGAAUAUUUUUGACUAAUCCCACUU<br>UACAUUAAUCUGUAUUGUGAAUGUGGAUAUUGAAUUAUA<br>UUUCUUUGUAAUCCCAUUAUCCAAAAUCCAGUUCAGAGAC<br>UAUUAGUUACCAAUGUUCACUGUGAAGGAAAAAAAAAAAA<br>AAAAAGCUCAGAGGAUAAACAUGUGAUAUGGUUUGGCUG<br>UGUCCCCACCCAAAUAUCAUCUUGAAUUGUAGCUCCCAU<br>AAUUCCCACGUGUUGUGGGAGGGACCCGGUGGGAGAUA<br>AUUGUAUCAUGGGGGUGGUUCCCCCAUACUAUUCUCAUA<br>GUAGUGAAUAAGUCUCACAAAAUCUGAUGGUUUUAUGAG<br>GGAAAACCCCUUUCACCUGGUUCUCAUUCUCUUCUCUGG<br>UCUGUCGUCAUGUAAGACAUGCCUUUCACCUUCUCCACC<br>AUGACUGUGAGGCCUCCCCAGCCACGUGGAACUGUGAGC<br>CCAUUAAACCUCUUUCACUUAUAAAU |
| Arm A | UGUCCUAUGAAUAUUUUUGACUAAUCCCACUUUACAUUAA<br>UCUGUAUUGUGAAUGUGGAUAUUGAAUUAUAUUUCUUUG<br>UAAUCCCAUUAUCCAAAAUCCAGUUCAGAGACUAUUAGUU |

| Sequence Name | Sequence (5'→3') |
|---|---|
| | ACCAAUGUUCACUGUGAAGGAAAAAAAAAAAAAAAAAGCU CAGAGGAUA |
| Arm B | AACAUGUGAUAUGGUUUGGCUGUGUCCCCACCCAAAUAU CAUCUUGAAUUGUAGCUCCCAUAAUUCCCACGUGUUGUG GGAGGGACCCGGUGGGAGAUAAUUGUAUCAUGGGGGUG GUUCCCCCAUACUAUUCUCAUAGUAGUGAAUAAGUCUCA CAAAAUCUGAUGGUUUUAUGAGGGAAAACCCCUUUCACC UGGUUCUCAUUCUCUUCUCUGGUCUGUCGUCAUGUAAGA CAUGCCUUUCACCUUCUCCACCAUGACUGUGAGGCCUCC CCAGCCACGUGGAACUGUGAGCCCAUUAAACCUCUUUCA C |
